# Structures of bacterial and human phosphoglycosyltransferases bound to a common inhibitor inform selective therapeutics

**DOI:** 10.64898/2025.12.16.694696

**Authors:** Beebee Yusrah Kaudeer, Jacob M. Kirsh, Katsuhiko Mitachi, Jessica M. Ochoa, Marie-Therese Soroush-Pejrimovsky, Yancheng E. Li, Vy N. Nguyen, Michio Kurosu, William M. Clemons

## Abstract

Glycoconjugates facilitate myriad biological processes, including cell–cell recognition and immune response, and they are generated by enzymes that transfer glycans. The orthologs MraY and DPAGT1 are dimeric phosphoglycosyltransferases involved in oligosaccharide biosynthesis for either bacterial peptidoglycan or eukaryotic *N*-linked glycans, respectively. Both enzymes play central regulatory roles, making them attractive targets for antibacterial and anticancer therapies. In our prior studies, a muraymycin A1-derived inhibitor termed APPB (aminouridyl phenoxypiperidinbenzyl butanamide) was developed. It exhibits sub-100 nM IC_50_ values against both MraY and DPAGT1 and has demonstrated efficacy against DPAGT1-dependent cancers, making it an excellent starting point for next-generation small molecules. To guide inhibitor development, we determined cryo-EM structures of APPB bound to MraY or DPAGT1 at 2.9 Å resolution using single-particle analysis. The structures reveal that APPB, composed of a nucleoside, a central amide, and a lipid-mimetic, adopts two conformations in each protein, which correlate with local hydrogen-bonding contacts of the central amide carbonyl. Examination of the amide carbonyl environments guides conformer selection for future DPAGT1-targeting anticancer agents. Further, comparisons of APPB-bound geometries and nucleoside interactions inform opportunities for antibacterial agents targeting MraY. Overall, our study provides design principles for MraY- or DPAGT1-specific drugs and motivates the utility of simultaneously characterizing inhibitor-bound orthologs for selective therapeutics.

---

Across all domains of life, glycans are appended to macromolecules to support essential biological functions. These glycans are typically synthesized on lipid carriers before being transferred to their final targets. In diverse pathways utilizing glycans, integral membrane phosphoglycosyltransferases of the polyprenyl-phosphate *N*-acetylhexosamine phosphate transferase (PNPT) superfamily play crucial roles.^1,2^ This is the case for two orthologous sub-100 kDa PNPTs, MraY (phospho-MurNAc pentapeptide translocase) in peptidoglycan biosynthesis and DPAGT1 (dolichyl-phosphate alpha-*N*-acetyl-glucosaminyl-phosphotransferase) in human *N*-linked glycosylation, which perform analogous reactions where a phosphosaccharide is transferred to a polyisoprenyl phospholipid.^3,4^ Each protein catalyzes a feedback-regulated step within interconnected glycosylation pathways, making them critical control points in their respective pathways.^5,6^ Microbes exploit this vulnerability by targeting other species’ PNPTs for their own survival: the bacteriophage ΦX174 single-gene lysis protein E inhibits *E. coli* MraY, causing host lysis, and *Streptomyces* spp. produce the secondary metabolite tunicamycin, a PNPT inhibitor that induces antibacterial activity against other prokaryotic organisms.^7–10^

The natural success of targeting PNPTs indicates that this approach could be a promising therapeutic strategy for human disease. The central role of MraY in peptidoglycan biosynthesis makes it an attractive target in the face of rising antibiotic resistance.^11^ In addition, many solid cancers are associated with aberrant (*N-*linked) glycosylation, and the upregulation of DPAGT1 is observed in many cancer cell lines.^12–15^ Treatment of cancer cells with tunicamycin *in vitro* has demonstrated antiproliferative effects, possibly due to its ability to inhibit DPAGT1.^16,17^ However, tunicamycin itself cannot be used as a therapeutic *in vivo*, as it has moderate IC_50_ values for MraY and DPAGT1 and exhibits severe toxicity towards healthy human cells.^17–20^ Nevertheless, tunicamycin-related molecules represent an important foundation for novel treatments, so long as improved analogs deliver stronger inhibition with selective toxicity.

With this hypothesis in mind, derivatives of muraymycin A1 – another *Streptomyces* antibiotic structurally similar to tunicamycin – were previously screened in search of strong DPAGT1 inhibitors.^21^ This process identified APPB (aminouridyl phenoxypiperidinbenzyl butanamide), which is comprised of a uridine nucleoside, a 5’-aminoribose, a central amide, a primary amide, and a TMPA (trifluoromethoxy phenoxy piperidin-1-yl phenyl methoxymethyl) lipid-mimetic tail (**Figure 1a**).^21^ APPB has significant potential as an anticancer therapeutic, with an IC_50_ of 25 nM for DPAGT1 and water solubility of ∼75 mg/mL.^15,21,22^ It has been highly efficacious in application: APPB induces apoptosis across diverse DPAGT1-dependent solid tumors without causing cytotoxicity, and it significantly reduces tumor growth in both xenograft and orthotopic mouse models.^19,22^ Further, prolonged dosing (10 mg/kg daily for 45 days) in mice produced no detectable changes in serum or plasma chemistry, highlighting its benign safety profile.^15,23^ In addition, APPB has an IC_50_ of 80 nM for MraY and exhibited potent bactericidal activity against spore-forming bacilli, notably *Clostridioides difficile*, even under strict anaerobic conditions and at low effective concentrations.^24^ As such, APPB is an excellent starting point for next-generation therapeutics for both DPAGT1 and MraY. However, structural information for APPB binding to either enzyme is unknown, thus, specific interaction-guided optimization is not yet possible. Therefore, we sought structures of drug-bound proteins to enable rational design. Pursuing both structures is beneficial for developing both MraY- and DPAGT1-selective therapeutics without cross-reactivity.^25,26^

**Figure 1.**
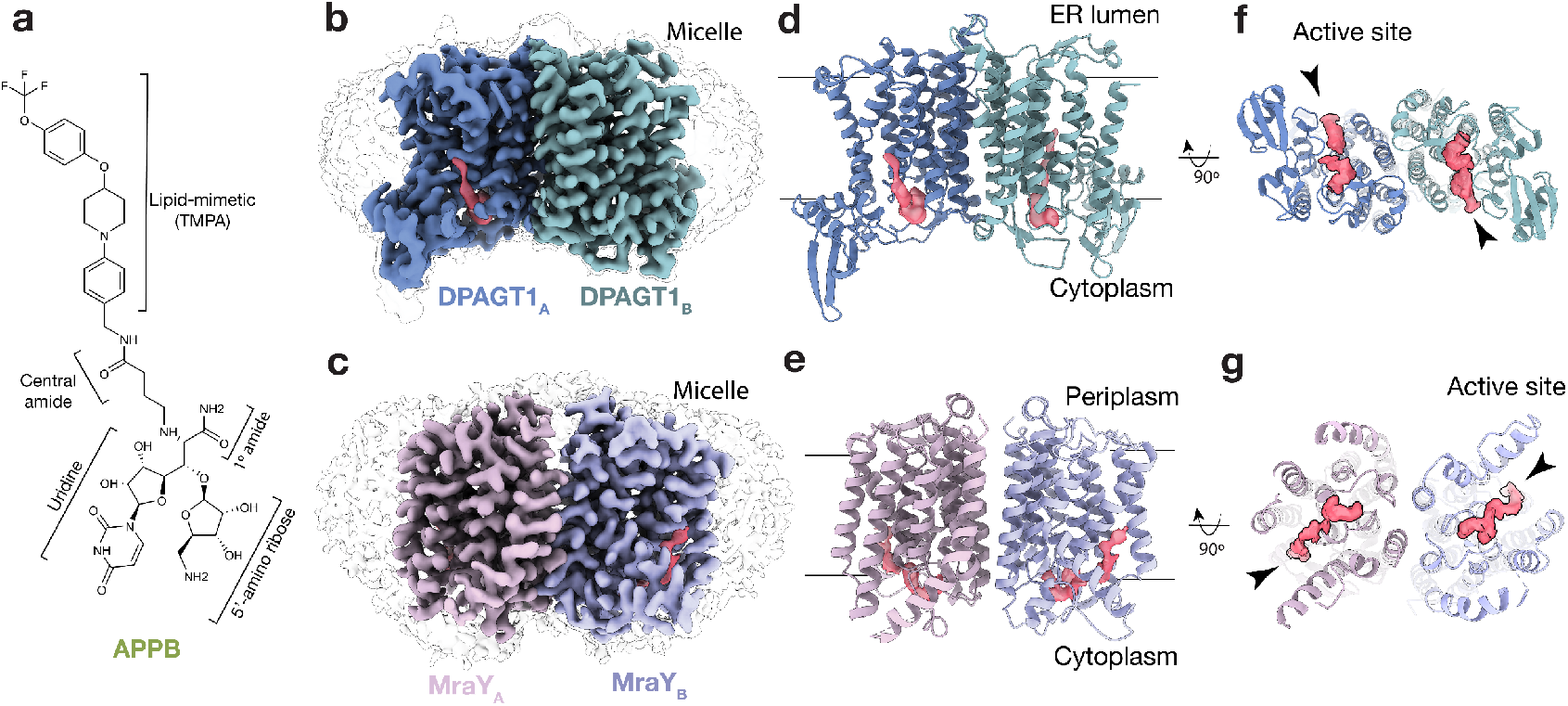
Cryo-EM maps show APPB binds at DPAGT1 and MraY active sites. (a) Chemical structure of APPB (aminouridyl phenoxypiperidinbenzyl butanamide), a potential antibacterial and anticancer drug comprised of a uracil nucleoside, a 5’-aminoribose, a central amide, a primary amide, and a TMPA (trifluoromethoxy phenoxy piperidin-1-yl phenyl methoxymethyl) lipid-mimetic tail.^21^ (b,c) Cryo-EM Coloumb potential maps (contoured at 0.07 in ChimeraX^27^) of DPAGT1 and MraY dimers bound to APPB viewed in the plane of the membrane (density corresponding to monomers are colored blue/green and pink /purple, respectively, with APPB density in red). A lower contour map in grey highlights the detergent micelles. (d,e) Cartoon models of DPAGT1 and MraY structures oriented and colored as in (b,c). Bars represent approximate boundaries of the membrane bilayer. (f,g) Cytoplasmic view of (d,e). Active sites and lipid-binding groove indicated by arrowheads.

## Hydrogenivirga sp

MraY and *H. sapiens* DPAGT1 were expressed and purified from *E. coli* using DM detergent and human-derived Expi293 cells using DDM detergent, respectively (**Figure S1**). Our purification of DPAGT1 from a human cell line differs from prior work, which expressed the protein in insect cells.^4,28^ Purified MraY and DPAGT1 were incubated with APPB (**Figure S2**) and vitrified for cryo-electron microscopy (cryo-EM) studies. Movies were acquired using a Titan Krios operating at 300 keV and 130,000x magnification. Coulomb potential maps of MraY and DPAGT1 bound to APPB were solved to a resolution of 2.9 Å using single-particle analysis (**Figure S3–S4; Table S1**). Importantly, these were solved without the addition of fiducial markers.

Previous structures of MraY and DPAGT1 indicated that they are homodimers, with monomeric units comprising 10 transmembrane helices (**Figure S5**).^4,28,29^ Overall, they have similar tertiary structures and active sites (**Figure S6**), but differ in their dimer interfaces.^28,30^ As expected, the cryo-EM maps reveal two monomers per protein, with residual density from detergent micelles and small molecules (**Figure 1b,c**). Dimeric proteins were modeled into the maps (**Figure 1d,e**). Interestingly, overlaying both DPAGT1 chains indicates symmetry breaking in select sidechain conformations (R40, K125, and Y256; **Figure S7**), whereas the MraY sidechains are superimposable within the limits of our resolution. Remaining ordered nonproteinogenic density was observed in each active site and is consistent with bound APPB, implying it binds in the same region as related orthosteric inhibitors (**Figure S8–S10**).^4,10,28,31^ A view from the cytoplasm highlights the distinct dimer interface and differing orientations of the active sites (**Figure 1f,g**).

For both orthologs, close inspection of the active site density reveals each has two distinct APPB conformations (**Figure 2a,b**). (Nearly) complete density is observed for all four conformers. In MraY, the density is consistent with both conformers being modeled in each active site with 50:50 occupancy (**Figure 2b**). For DPAGT1, the binding is asymmetric, where distinct conformers were modeled into each monomer (**Figure 2a**). This observation mirrors the symmetry breaking observed in the DPAGT1 sidechains between the monomers (**Figure S7**). The modeling indicates that the APPB conformers are *cis*and *trans* isomers of the central amide; hereafter, the four conformers of APPB are referred to as *cis*_DPAGT1_, *trans*_DPAGT1_, *cis*_MraY_, or *trans*_MraY_ (**Figure 2a,b**). Overall, the APPB conformers in DPAGT1 have similar overall geometries (**Figure 2a**), while the MraY conformers have distinct TMPA lipid-mimetic geometries (**Figure 2b**).

**Figure 2.**
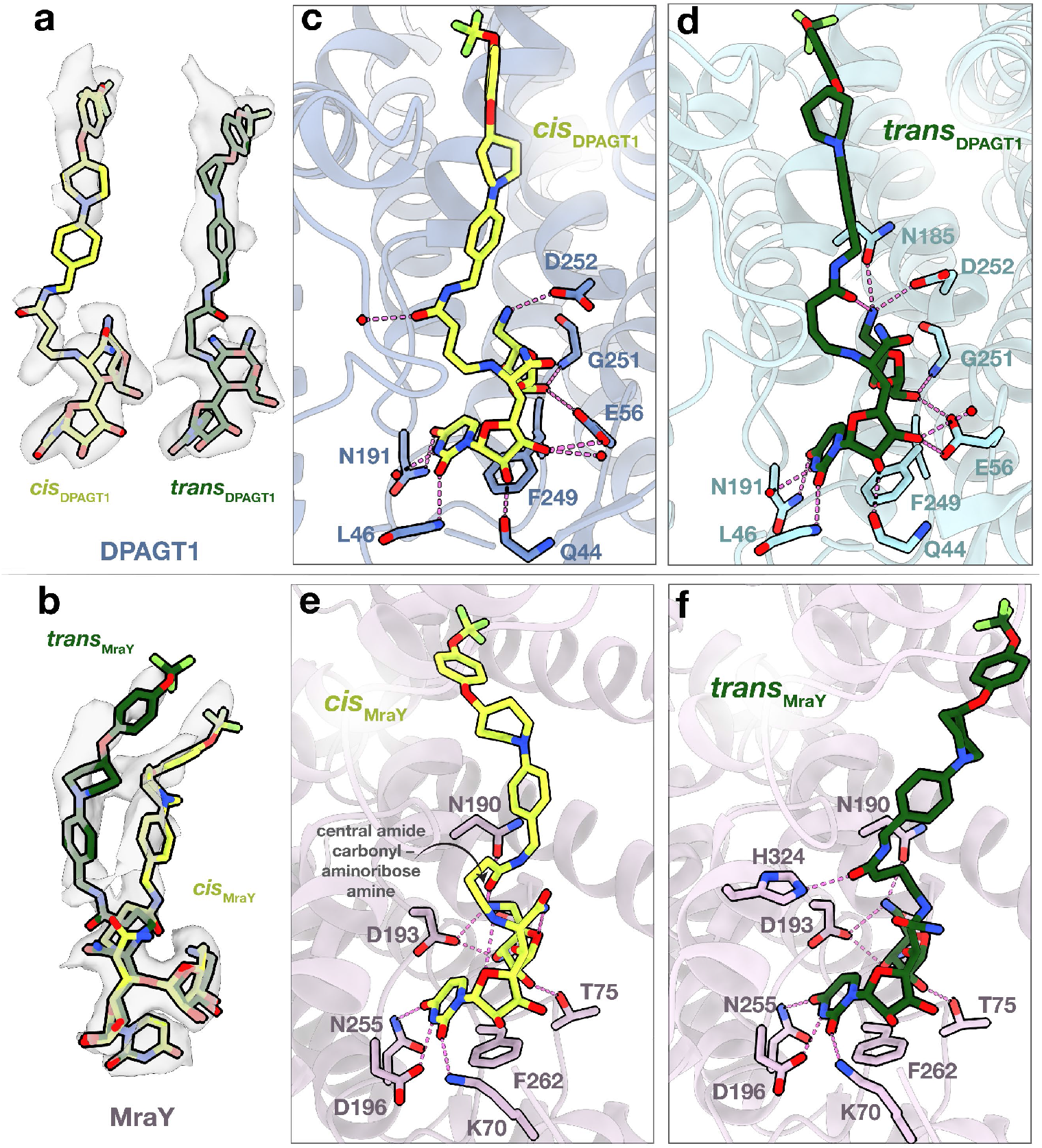
Interactions between DPAGT1/MraY and APPB. (a,b) Cryo-EM Coulomb potential maps of APPB in DPAGT1 (a) and MraY (b), with APPB modeled as sticks. APPB conformations correspond with *cis* (neon green) and *trans* (forest green) central amides. Conformers are referred to as *cis*_DPAGT1_, *trans*_DPAGT1_, *cis*_MraY_, or *trans*_MraY_. Detailed local interactions of *cis*_DPAGT1_ (c), *trans*_DPAGT1_ (d), *cis* _MraY_ (e), *trans*_MraY_(f). The protein is shown in ribbon form, APPB-contacting residues are shown as sticks, and waters are represented by red spheres. Dashed lines indicate potential H-bonding interactions based on non-hydrogen atom distances <3.5 Å. Schematic representations of APPB interactions can be found in **Figure S11**.

We inspected the local environments of APPB in DPAGT1 and MraY to inform APPB-based drug design. We infer potential hydrogen bonds (H-bonds) based on distances <3.5 Å between non-hydrogen atoms. In DPAGT1, the central amide carbonyl in *cis*_DPAGT1_ makes an *inter*molecular H-bond with a bound water (**Figure 2c**). In contrast, the carbonyl in *trans*_DPAGT1_ forms an *intra*molecular H-bond with the aminoribose amine, implying APPB geometry depends on the local H-bonding environment of the central amide carbonyl (**Figure 2c,d**). For the aminoribose amine, additional H-bonds are formed with N185 and D252 in *trans*_DPAGT1_, whereas the amine only H-bonds with D252 in *cis*_DPAGT1_ (**Figure 2c,d**). The aminoribose interactions are completed by H-bonds between the 3’-hydroxyl with the G251 backbone N–H and E56 sidechain in both conformers (**Figure 2c,d**). APPB uridine interactions are the same between the conformers and are similar to those seen in prior structures with the uridines in tunicamycin and UDP-GlcNAc (**Figure 2c,d,S8**).^4,28^

In MraY, analogous APPB interactions are observed as in DPAGT1, but with a key difference: the central amide carbonyl in *cis*_MraY_ forms an *intra*molecular H-bond with the aminoribose amine (**Figure 2e**), whereas the carbonyl makes an *inter*molecular H-bond with His324 in *trans*_MraY_ (**Figure 2f,S12**). In both conformers, the aminoribose amine H-bonds with the N190 and D193 sidechains, the 3’-hydroxyl H-bonds with the D193 sidechain, and the 2’-hydroxyl H-bonds with the T75 sidechain (**Figure 2e,f**). As in DPAGT1, the uridine binding is highly similar between the conformers and for MraY bound to various nucleoside inhibitors (**Figure 2e,f,S9–S10**).^10,31–34^ In contrast to DPAGT1, other inhibitors with the aminoribose moiety have been solved, and the interactions for MraY with this group are similar to those in muraymycin D2, carbacaprazamycinm, sphaerimicin analogue, and analogue 3 (**Figure S10**).^31–34^ Interestingly, no clear interactions are observed for the uridine ribose hydroxyls, whereas H-bonds are formed with both the 2’- and 3’-hydroxyls in DPAGT1 (**Figure 2c–f,S11**). Consequently, there are fewer APPB–protein contacts in MraY than in DPAGT1, which may account for the lower APPB IC_50_ for DPAGT1 than for MraY.

To uncover principles for selective inhibitors, we first considered APPB binding within DPAGT1 for the design of potential therapeutics such as anticancer drugs. In DPAGT1, the conformations have similar geometries (**Figure 2a**) but differ mainly in the orientation and local interactions of the central amide carbonyl (**Figure 2c,d**). As previously mentioned, the aminoribose amine in *trans*_DPAGT1_ H-bonds with the central amide carbonyl and two other H-bond partners to fulfill its H-bonding potential (**Figure 2d,S13**). In contrast, the amine in *cis*_DPAGT1_ does not H-bond with the carbonyl and only adopts one H-bond (**Figure 2c,S13**). The additional two H-bonds in *trans*_DPAGT1_ suggests this isomer may be optimized for DPAGT1-selective anticancer agents. Further derivatization of the primary amide, which H-bonds with water in both *trans*_DPAGT1_ and *cis*_DPAGT1_ (**Figure S11**), may help reinforce this binding geometry.

In light of its bifunctional therapeutic potential, we compared how APPB binds to MraY and DPAGT1 (**Figure 3**). We found that MraY has a wider lipid-binding groove than DPAGT1. This topological difference likely explains why APPB shows greater geometric differences between *cis*_MraY_ and *trans*_MraY_ compared to *cis*_DPAGT1_ and *trans*_D-PAGT1_ (**Figure 2**). It also suggests rational design strategies: the TMPA lipid tail could be made bulkier, so it no longer fits into the DPAGT1 lipid-binding groove. Additionally, the nucleoside ribose currently does not interact with MraY (**Figure 2e,f**) but does interact with DPAGT1 (**Figure 2c,e**). This suggests further APPB derivatization could introduce MraY-specific contacts while reducing DPAGT1 affinity.^26,35^ Finally, APPB shows different interactions with DPAGT1 and MraY conserved active site residues, and these differences may be exploited for ortholog specificity (**Figure S6**). These collective insights stem from our ability to resolve high-resolution membrane protein features, underscoring cryo-EM’s potential to facilitate therapeutic advances.^36^

**Figure 3.**
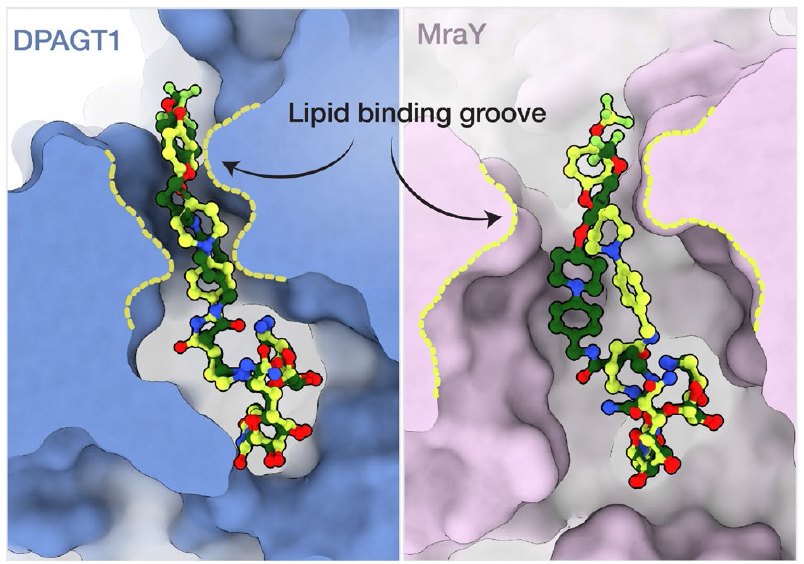
DPAGT1 and MraY structural comparison suggests differentiation strategies for antibiotics. Molecular surface views of the binding grooves of DPAGT1 (left) and MraY (right) with the *cis* and *trans* APPB conformers shown in sticks and colored as in **Figure 2**. DPAGT1 has a tighter lipid-binding groove compared to MraY, as indicated by the dashed yellow lines.

In summary, we report cryo-EM structures of PNPT family members DPAGT1 and MraY bound to APPB, a recently discovered small molecule with promising anticancer and antibacterial potential.^15,21^ Our results indicate that APPB adopts two distinct conformations in both orthologs, and direct comparison of its binding environment reveals chemically addressable opportunities for differentiation. These comparisons emphasize the power of obtaining structures of both human and bacterial orthologs to develop more selective inhibitors.^37–40^ Modern machine-learning-based techniques use chemical fragments to target specific binding-pocket regions for drug discovery, and these can assist with implementing our findings to develop next-generation APPBs.^41,42^ In conclusion, we reiterate that DPAGT1 and MraY are compelling targets due to their central roles in conserved pathways. Yet, key questions remain about the conserved chemical and catalytic mechanism of the PNPT superfamily. As mechanistic insights expand regulatory opportunities, future work will leverage the structural pipelines developed herein to enable more efficacious therapeutics.

## ASSOCIATED CONTENT

### Supporting Information

The Supporting Information is available free of charge on the ACS Publications website.

Expression, purification, cryo-EM sample and grid preparation, data collection and processing, and model building for DPAGT1 and MraY, DPAGT1 and MraY genes, APPB HPLC analysis, DPAGT1 and MraY sequence and structural alignments, DPAGT1 model symmetry breaking, structural comparisons of DPAGT1 and MraY nucleoside inhibitors, contact maps for APPB–protein interactions, comparison of APPB intramolecular and intermolecular H-bonding interactions, comparison of H-bonds formed by APPB aminoribose amine in *trans*_DPAGT1_ and *cis*_DPAGT1_, and cryo-EM data collection, processing, refinement, and validation statistics.

## Author Contributions

M.K. and W.M.C. conceptualized the project. K.M. synthesized and purified APPB. J.M.O. and V.N.N. designed DPAGT1 expression construct, J.M.K., and M.-T.S.-P. expressed and purified DPAGT1, B.Y.K., J.M.K., M.-T.S.-P., and Y.E.L. collected and processed cryo-EM data, and J.M.K. performed model building, refinement, and validation of DPAGT1. B.Y.K. performed all analogous tasks for MraY. B.Y.K., J.M.K., and W.M.C. wrote the manuscript with input from all authors. All authors have given approval to the final version of the manuscript.

## ACKNOWLEDGMENT

Funding for this work was provided by National Institutes of Health grant R01GM114611 (W.M.C. and M.K.), the G. Harold and Leila Y. Mathers Foundation (W.M.C.), and Biohub (W.M.C). Cryo-EM data was performed in the Beckman Institute Resource Center for Transmission Electron Microscopy at Caltech, and we thank Songye Chen for assistance with data collection. We thank Rebecca Voorhees for support and kindly providing materials necessary for DPAGT1 expression and purification. We thank Grace Baron for feedback on the manuscript and Andrew J. Saks for assistance with processing DPAGT1 cryo-EM data.

## ABBREVIATIONS

PNPT: polyprenyl-phosphate *N*-acetylhexosamine phosphate transferase
MraY: phospho-N-acetylmuramoyl-pentapeptide-transferase
DPAGT1: dolichyl-phosphate alpha-*N*-acetyl-glucosaminyl-phosphotransferase
APPB: aminouridyl phenoxypiperidinbenzyl butanamide
TMPA: trifluoromethoxy phenoxy piperidin-1-yl phenyl methoxymethyl
cryo-EM: cryo-electron microscopy
DM: n-Decyl-β-Maltoside
DDM: n-Dodecyl-β-Maltoside.

## ACCESSION CODES

APPB-bound DPAGT1: 9ZNN and EMD-74451

APPB-bound MraY: 9ZNO and EMD-74452

## Insert Table of Contents artwork here

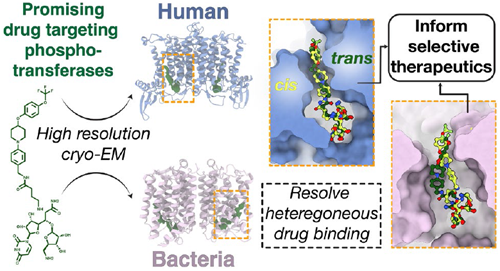

