## Supplemental Information for "Structures of bacterial and human phosphoglycosyltransferases bound to a common inhibitor inform selective therapeutics"

### Methods

#### DPAGT1 expression and purification

Plasmid construction

A *Homo sapiens* DPAGT1 gene (*Hs*DPAGT1; UniProtKB Q9H3H5) was designed and purchased from Twist Biosciences (San Francisco, CA). The gene was cloned into a pHAGE2 backbone downstream of a CMV promoter and upstream of a TEV cleavage site, a GFP gene, a P2A ribosomal skip site,[1] and an mCherry reporter gene.[2] The construct was designed so for three reasons: lentiviral transduction efficacy can be assessed by monitoring mCherry signal, proper DPAGT1 membrane localization can be assessed by monitoring GFP signal, and recombinant DPAGT1-GFP can be purified via GFP-immunoprecipitation (GFP-IP).[3]

Lentiviral transduction

Human embryonic kidney derived adherent 293T (HEK293T; ATCC CRL-11268) cells were thawed and resuspended in DMEM (Lonza 12-614Q) media supplemented with 2 mM glutamine and 10% (w/v) fetal bovine serum (FBS; Thermo Fisher A3382001). Cells were pelleted at 300*g* for 5 minutes and distributed into a 15-cm plate containing the supplemented DMEM media and incubated at 37˚C in 5% $CO_2$ for 16 hours. Once cells were approximately 80–90% confluent, media was aspirated, and cells were washed with 4 mL of DPBS (Thermo Fisher 14040117). 4 mL of 0.25% Trypsin (Thermo Fisher 25200-056) was added and incubated for 2 minutes to detach cells. The reaction was quenched by adding 16 mL of DMEM to the plate. The cells were counted, and 750,000 cells were transferred to each well in a 6-well plate to grow overnight to perform 2.5 mL transfection reactions. The following day, when 80–90% confluency was reached, cells were incubated with a mixture of 0.25 mL Opti-MEM (Gibco2631985-062), 0.94 µg psPAX2 2$^{nd}$ generation lentiviral packaging plasmid (Addgene), 0.32 µg pMD2.G lentiviral envelope plasmid (Addgene), 1.25 µg of pHAGE2 plasmid, and 7.5 µL TransIT-293 transfection reagent (Mirus MIR2704) according to the manufacturer's instruction. After 8 hours, media was exchanged with fresh DMEM, and cells were incubated for approximately 16 hours at 37˚C in 5% $CO_2$. Media containing lentiviral particles (serum) was harvested, flash frozen, and stored at -80˚C. On the day of lentiviral transduction, 10 x $10^6$ Expi293F cells (Thermo Fisher A14527) at greater than 95% viability were incubated with 3 mL of thawed lentiviral serum and Polybrene Transfection Reagent (8 µg/mL final concentration; Millipore Sigma TR-1003-G) in a shaking incubator at 37˚C with 8% $CO_2$. After 8 hours, media was exchanged, and cultures were maintained between 0.5 x $10^6$ and 2 x $10^6$ cells/mL. Transduction success and DPAGT1 expression were monitored by mCherry and GFP signals, respectively, using an EVOS M5000 microscope (Thermo Fisher). HEK293T and Expi293F cells and lentiviral packaging and envelope plasmids were kind gifts from the Voorhees lab (Caltech).

Fluorescence activated cell sorting

Expi293F cells expressing DPAGT1 were sorted on a SONY SH800S (Sony Biotechnology) using fluorescence activated cell sorting (FACS) by monitoring GFP signal at 488 nm. 1.5 x $10^6$ transduced Expi293F cells were sorted into a six-well plate, which was incubated at 37 °C and 8% $CO_2$ for 3 days with shaking at 120 rpm. Cells were transferred to 20 mL of fresh Expi293 Expression Medium (Thermo Fisher A1435101) in a 125-mL vented Erlenmeyer flask. When the cell density reached 2 x $10^6$ cells/mL, cells were transferred to 100 mL of fresh Expi293 media in a 1000-mL vented flask. Once cells were reliably doubling every 24 hours, they were harvested, frozen, and maintained as 15 x $10^6$ cells/mL aliquots under cryogenic conditions in Expi293 media supplemented with 10% DMSO. Aliquots were thawed, passaged, and maintained as needed.

DPAGT1-GFP expression & GFP-IP purification

Expi293F cells harboring the pHAGE2 plasmid were grown and maintained using Expi293 media in a humidified 8% $CO_2$ incubator on an orbital shaker at 125 rpm. 400 – 450 mL of cell culture were grown to a density of 8 – 12 x $10^6$ cells/mL. Cells were harvested at 4,500*g* for 15 minutes at 4˚C. Supernatant was removed and the cells were resuspended in 20 mL of cold DPBS and pelleted at 3,200*g* for 15 minutes at 4˚C. Supernatant was removed and cell pellets were weighed and used immediately for purification or frozen and stored at -80°C. All subsequent purification steps were performed at 4˚C.

GFP-IP purification closely followed the method in ref. [3]. GFP-IP requires biotinylated anti-GFP nanobody harboring a $SUMO^{Eu1}$ tag and cognate $SENP^{EuB}$ protease.[4] The nanobody and protease are encoded by *E. coli* expression vectors pTP396 (Addgene ID 149336) and pAV0286 (Addgene ID 149333), respectively. The nanobody and protease were expressed in AVB101 (Avidity CVB101) and NEBExpress (New England Biolabs E5360) chemically competent cells, respectively, as previously described.[3] Both plasmids were kindly provided by the Voorhees lab (Caltech).

Cells pellets were resuspended in a solubilization buffer containing 50 mM HEPES pH 7.5, 200 mM NaCl, 5 mM $MgCl_2$, 1% (w/v) n-dodecyl-β-D-maltopyranoside (DDM, Anatrace), protease inhibitor cocktail (Roche 11836153001), and 2 mM TCEP (UBPBio P1020). To avoid excessive dilution, 6.8 mL of solubilization buffer were used per 1 g of cell pellet. Cells were fully resuspended by vortexing and solubilized for 1.5 hours at 4˚C with gentle rocking. Following solubilization, large debris and aggregates were removed by centrifugation at either 34,500*g* for 1 hour or 39,000*g* for 35 minutes at 4˚C. Simultaneously, anti-GFP nanobody was immobilized on magnetic Streptavidin beads (Thermo Fisher PI88817) by shaking on a thermomixer (Eppendorf Thermomixer R) at 1300 rpm and 4˚C. For simplicity, this anti-GFP nanobody:magnetic Streptavidin bead conjugate will be referred to as *magnetic SA beads*. The supernatant containing solubilized DPAGT1-GFP was incubated with the magnetic SA beads for 1.5 hours at 4˚C with gentle rocking. The supernatant and unbound debris were removed, and magnetic SA beads were washed with a wash buffer containing 50 mM HEPES pH 7.5, 200 mM NaCl, 5 mM $MgCl_2$, 0.022% (w/v) DDM, and 2 mM TCEP. The magnetic SA beads were incubated with $SENP^{EuB}$ protease for 1 hour at 4˚C on the thermomixer shaking at 1300 rpm to elute DPAGT1-GFP. The supernatant was collected, and the magnetic SA beads were discarded. As a modification to the protocol in ref. [3], TEV protease (New England Biolabs P8112S) was added to the supernatant at a ratio of 1 µL TEV protease : 20 µg protein. Cleavage of DPAGT1 and GFP was allowed to proceed overnight at 4˚C with shaking at 1300 rpm on the thermomixer. The supernatant was collected, concentrated to ~500 µL with a 100 kDa MWCO Centrifugal filter (Amicon), and DPAGT1 was isolated using size exclusion chromatography on a Superdex 200 Increase 10/300 GL size exclusion chromatography column (Cytiva) pre-equilibrated with wash buffer (**Figure S1**). Analysis of the chromatogram indicated >90% cleavage efficiency. Fractions were analyzed by SDS-PAGE.

**MraY expression and purification**

NiCo21 (DE3) pLEMO (DE3) competent cells were transformed with pET22b-His6-*Hy*MraY plated in LB-agar containing 34 µg/ml chloramphenicol and 50 µg/mL ampicillin. Colonies from the plate were scraped off and resuspended in 30 mL of 2x Yeast Extract Tryptone (2xYT) media to inoculate 6L of 2xYT with 34 µg/ml chloramphenicol, 50 µg/mL ampicillin, and 0.4 mM L-Rhamnose. Cells were grown shaking at 225 rpm at 37 °C and were transferred to 30˚C when optical density ($OD_{600}$) reached 0.6 – 0.7. Expression was induced with the addition of 400 µM Isopropyl β-D-1-thiogalactopyranoside (IPTG). Culture was harvested after approximately 4 hours by centrifugation at 9,000*g* for 10 minutes at 4 ˚C. Cell pellets were frozen and stored at -80°C or used immediately for purification. All subsequent purification steps were performed at 4˚C.

Cell pellets were resuspended in lysis buffer (20 mM Tris pH 7.5, 300 mM NaCl, 10% glycerol, 5 mM β-mercaptoethanol (βME), and protease inhibitors: 1 mM phenylmethylsulfonyl fluoride (PMSF) and benzamidine (Benz)). Re-suspended cells were homogenized and lysed through four passes using a M-110L microfluidizer (Microfluidics). Unlysed cells and cell debris were removed by centrifugation at 23,500*g* for 30 minutes. The supernatant was then centrifuged at 168,000*g* for 1 hour to obtain the cell membrane. Membrane pellets were flash frozen in liquid nitrogen and stored at -80°C.

The membrane pellet was homogenized and solubilized for 2 hours in extraction buffer (20 mM HEPES pH 7.5, 300 mM NaCl, 10% glycerol, 1% (w/v) n-Decyl-β-Maltoside (DM, Anatrace) and 10 mM imidazole). The membrane extraction was centrifuged at 168,000*g* for 30 minutes to remove unextracted membrane proteins. The supernatant was incubated with 1 mL of Ni-NTA agarose resin (Qiagen) equilibrated with extraction buffer for 1 hour. The incubated sample was loaded onto a gravity column and washed with 50 column-volumes of wash buffer A (10 mM HEPES pH 7.5, 300 mM NaCl, 5% glycerol, 0.15% DM and 20 mM imidazole) and another 50 column volumes of wash buffer B (10 mM HEPES pH 7.5, 300 mM NaCl, 5% glycerol, 0.15% DM and 20 mM EDTA). The protein was eluted with 5 column-volumes of elution buffer (20 mM HEPES pH 7.5, 150 mM NaCl, 5% glycerol, 0.15% DM and 200 mM EDTA). Eluted protein was concentrated with a 100 kDa MWCO filter to ~500 µL and run through a Superdex 200 Increase 10/300 GL size exclusion chromatography column pre-equilibrated with 20 mM HEPES pH 7.5, 75 mM NaCl, 5% glycerol, 0.15% DM and 10 mM $MgCl_2$. Fractions were analyzed by SDS-PAGE.

**Cryo-EM sample and grid preparation**

Purified MraY and DPAGT1 were separately pooled and concentrated with a 100 kDa MWCO filter to ~12 mg/mL (MraY) and ~4 mg/mL (DPAGT1) for cryo-EM grids. 0.4 mM of APPB (aminouridyl phenoxypiperidinbenzyl butanamide)-HCl in 20 mM HEPES pH 7.5, 100 mM NaCl was added to the sample and incubated on ice for at least 1 hour. 0.05% of 3-[(3-Cholamidopropyl)dimethylammonio]-2-hydroxy-1-propanesulfonate (CHAPSO, Anatrace) was added to sample immediately before freezing. 3 µl of sample was applied to Quantifoil holey carbon R1.2/1.3 300 Mesh, Copper (Quantifoil, Micro Tools GmbH) grids that were glow-discharged for 75 seconds with a current of 20 mA at a pressure of 0.30 bars. The grids were blotted with an FEI Vitrobot Mark IV at 4°C, 100% humidity for 3 seconds with a blot force of +8 and immediately plunged into liquid ethane.

**Cryo-EM data collection and image processing**

Grids were screened for ice thickness and sample quality using a 200 keV Talos Arctica TEM equipped with a Gatan K3 direct electron detector. Cryo-EM data was acquired on a Thermo Fisher Scientific Titan Krios operating at an acceleration voltage of 300 keV and equipped with a Gatan Energy Filter (20 eV slit width) and a Gatan K3 detector operating in super-resolution mode. All movie stacks were acquired with a defocus range of −1.0 to −3.0 µm at a nominal magnification of 130,000x, corresponding to a raw pixel size of 0.325 Å/pixel. 40-frame movies were recorded using an automated acquisition pipeline in SerialEM[5] and were subjected to a total dosage of 70 $e^-/Å^2$. Correlated double sampling (CDS) mode was enabled to improve image signal-to-noise ratio.[6]

Data collected from the Krios were processed in cryoSPARC (v4.7.1).[7] Movies were motion-corrected using "Patch Motion Correction" with a Fourier-crop of ½, resulting in a pixel size of 0.65 Å/pixel. Contrast transfer function (CTF) parameters for the resulting micrographs were estimated with "Patch CTF Estimation". Micrographs were manually curated based on CTF fits (≤4 Å), total frame motion, and relative ice thickness.

For DPAGT1, particles were selected by "Blob Picking" and extracted with a 480-pixel box size and a 2x bin (1.30 Å/pixel). The resulting particles were cleaned through iterative 2D classification to retain high-quality particle selections. 180,879 particles were used for "Ab-Initio Reconstruction" to generate three initial 3D models, and one model showed clear density for transmembrane domains. The corresponding 106,099 particles were run through "Non-Uniform Refinement"[8], resulting in an initial 3.5 Å reconstruction. The particles were used to train a Topaz[9] model to find high-quality particles not selected through blob picking, and the particles were again cleaned through iterative 2D classification. The remaining particles were combined with the high-quality particles from blob picking, and duplicates were removed. The combined particles were iteratively classified and filtered with ab-initio reconstruction to eliminate poorly resolved classes. A volume with clear transmembrane domains was subjected to successive "Global CTF Refinement" or "Local CTF Refinement" jobs followed by non-uniform refinement until the map resolution and quality no longer improved. To further improve map quality, "Reference-Based Motion Correction" was run with a 1x bin (0.65 Å/pixel) followed by additional global/local CTF refinement and non-uniform refinement jobs. Early attempts at model building into the resulting map identified several symmetry breaking sites between DPAGT1 protomers (**Figure S7**), so a final non-uniform refinement on 82,209 particles was run with relaxed C2 symmetry (maximization method). This yielded a 2.9 Å map with better overall quality than the corresponding C1 map. For all non-uniform refinement jobs, the overall resolution was estimated from the gold-standard Fourier shell correlation (GSFSC) curve at a cut-off of 0.143. After initial model docking and refinement (see below), masks were created for chains A and B, and these were used for "Local Refinements". The two locally refined maps were used to generate a composite map in ChimeraX[10] as described in refs. [11,12]. Local resolution of the symmetry relaxed map was obtained using the "Local Resolution Estimation" job, while local resolution of the composite map (shown in **Figure S3f**) was calculated by running the job for each of the locally refined maps. A detailed flowchart of DPAGT1 data processing is presented in **Figure S3**. The unsharpened, sharpened (B-factor: -79.7 $Å^2$), and locally refined maps were exported for model building, as well as a map from the "Local Filtering" job that filtered the unsharpened map using its local resolution estimation.

For MraY, good particles were picked with a Topaz[9] model trained on a previously collected, unpublished MraY dataset, extracted with a 480-pixel box size and a 2x bin (1.30 Å/pixel), and cleaned through iterative 2D classification. The high-quality particles were subjected to ab-initio reconstruction to generate initial 3D models and to eliminate poorly resolved classes. A subset of 154,920 particles yielding maps with well-defined transmembrane domains were selected to perform multiple rounds of 3D classification using two-class ab-initio reconstruction. The highest quality particles were run through non-uniform refinement, resulting in an initial 3.2 Å reconstruction. To further improve map quality, reference-based motion correction was run on the remaining 107,145 particles with a 1x bin (0.65 Å/pixel) followed by additional global CTF refinement, local CTF refinement, and non-uniform refinement jobs. The 3D reconstruction was analyzed with an early model built to check for symmetry. Unlike DPAGT1, no obvious symmetry breaking was observed, so C2 symmetry was applied to a final round of non-unform refinement, yielding a 2.9 Å map (note that no symmetry was applied until this point to ensure no bias was introduced). The overall resolution for MraY was estimated from the gold-standard Fourier shell correlation (GSFSC) curve. Both unsharpened and sharpened maps (B-factor: -109.8 $Å^2$) were exported from cryoSPARC for model building and refinement. The local resolution of the map was calculated using local resolution estimation job. A detailed flowchart of MraY data processing is presented in **Figure S4**.

##### DPAGT1 model building and refinement

The unsharpened map was run through the EMReady[13] web server (http://huanglab.phys.hust.edu.cn/emready/) to improve interpretability of (low-resolution) map features. An AlphaFold2[14] prediction of *Hs*DPAGT1 was docked into the EMReady map as a rigid body in UCSF ChimeraX[10]. In Coot[15] (v.0.9.8), manual refinements were made to the model such that all residues fit well in the EMReady map, followed by real-space refinement in Phenix[16] (v1.21.2-5419). CIF restraints for the APPB ligand were generated on Grade2 Web Server (https://grade.globalphasing.org/)[18]. APPB was manually fitted into the residual, nonproteinogenic density of the unsharpened map, and model manual refinement in Coot followed by refinement in Phenix was performed. Ordered waters near the active site were docked into density in the sharpened map, again followed by manual refinement in Coot and refinement in Phenix. Manual refinement in Coot and refinement in Phenix was also performed using the local filtering map. Manual adjustments were made to Chain A or B and associated ligands in Coot against the corresponding local refinement maps. These maps and model were used to generate the composite map,[11,12] and the model was refined against the composite map using real-space refinement in Phenix. Residues which were absent in the density or highly disordered (parts of the N- and C-termini and residues 80–89) were removed from the model. The final structural model was validated in Molprobity[17].

##### MraY model building and refinement

An AlphaFold2[14] prediction of dimeric *Hy*MraY was docked into the unsharpened map as a rigid body in UCSF ChimeraX[10]. The model was built manually into good quality regions of the unsharpened map in Coot[15] (v.0.9.6). Density for Loop 1 (residues 56–66) was absent, consistent with the crystal structure and likely reflecting high flexibility in this region. Through iterative rounds of manual refinements, the initial model was refined in Coot and in Phenix[16] using real-space refinement. APPB was manually fitted into the residual, nonproteinogenic density of the sharpened map, and another round of manual refinement in Coot and Phenix refinement was performed. The final model was visually inspected and validated on Molprobity[17]. In this model, two APPB conformers are present in each active site with 50:50 occupancies.

**Statistics and Availability for Maps & Models**

Cryo-EM data collection, processing, refinement, and validation statistics are presented in **Table S1.** Structures have been deposited into the PDB and Electron Microscopy Data Bank (EMDB) databases with accession codes 9ZNN/EMD-74451 and 9ZNO/EMD-74452 for DPAGT1 and MraY, respectively.

#### Genes

##### ***Hs*DPAGT1-TEV-GFP-P2A-RFP**

DPAGT1, GFP, and RFP are colored blue, green, and red, respectively. DPAGT1-GFP construct which is expressed shown with capitalized letters. The TEV cleavage site is underlined and the P2A skip site is bolded.

ATGTGGGCCTTCTCGGAATTGCCCATGCCGCTGCTGATCAATTTGATCGTCTCGCTGCT
GGGATTTGTGGCCACAGTCACCCTCATCCCGGCCTTCCGGGGCCACTTCATTGCTGCG
CGCCTCTGTGGTCAGGACCTCAACAAAACCAGCCGACAGCAGATCCCAGAATCCCA
GGGAGTGATCAGCGGTGCTGTTTTCCTTATCATCCTCTTCTGCTTCATCCCTTTCCCCT
TCCTGAACTGCTTTGTGAAGGAGCAGTGTAAGGCATTCCCCCACCATGAATTTGTGGC
CCTGATAGGTGCCCTCCTTGCCATCTGCTGCATGATCTTCCTGGGCTTTGCGGATGATG
TACTGAATCTGCGCTGGCGCCATAAGCTGCTGCTACCTACAGCTGCCTCACTACCTCT
CCTCATGGTCTATTTCACCAACTTTGGCAACACGACCATTGTGGTGCCCAAGCCCTTC
CGCCCGATACTTGGCCTGCATCTGGACTTGGGAATCCTGTACTATGTCTACATGGGGCT
GCTGGCAGTGTTCTGTACCAATGCCATCAATATCCTAGCAGGAATTAACGGCCTAGAG
GCTGGCCAGTCACTAGTCATTTCTGCTTCCATCATTGTCTTCAACCTGGTAGAGTTGG
AAGGTGATTGTCGGGATGATCATGTCTTTTCCCTCTACTTCATGATACCCTTTTTTTTCA
CCACTTTGGGATTGCTCTACCACAACTGGTACCCATCACGGGTGTTTGTGGGAGATAC
CTTCTGTTACTTTGCTGGCATGACCTTTGCCGTGGTGGGCATCTTGGGACACTTCAGC
AAGACCATGCTACTATTCTTCATGCCCCAGGTGTTCAACTTCCTCTACTCACTGCCTCA
GCTCCTGCATATCATCCCCTGCCCTCGCCACCGCATACCCAGACTCAATATCAAGACA
GGCAAACTGGAGATGAGCTATTCCAAGTTCAAGACCAAGAGCCTCTCTTTCTTGGGC
ACCTTTATTTTAAAGGTGGCAGAGAGCCTCCAGCTGGTGACAGTACACCAGAGTGAG
ACTGAAGATGGTGAATTCACTGAATGTAACAACATGACCCTCATCAACTTGCTACTTA
AAGTCCTTGGGCCCATACATGAGAGAAACCTCACATTGCTCCTGCTGCTGCTGCAGAT
CCTGGGCAGTGCCATCACCTTCTCCATTCGATATCAGCTCGTTCGACTCTTCTATGATG
TCGGTTCGAGCGCTAGC<u>GAGAACCTGTATTTTCAGGGT</u>GTGTCTAAAGGTGAAGAAC
TTTTTACGGGTGTCGTCCCCATCCTGGTTGAATTGGACGGAGACGTGAATGGCCATAA
ATTTTCAGTGAGCGGTGAAGGAGAGGGAGACGCCACTTACGGTAAACTTACCTTGAA
ATTTATCTGCACTACCGGGAAACTGCCCGTACCCTGGCCGACGCTCGTAACAACGCTT
ACCTACGGCGTGCAATGTTTTAGCAGATACCCAGACCATATGAAGCAGCACGATTTTT
TCAAGAGTGCAATGCCTGAGGGGTACGTACAGGAAAGGACGATCTTTTTCAAAGACG
ATGGTAATTATAAAACGCGGGCTGAGGTTAAATTTGAAGGAGACACGCTCGTGAATC
GAATAGAGCTGAAGGGAATCGACTTCAAGGAAGACGGTAACATCCTGGGCCATAAAC
TCGAATATAATTACAATTCACACAACGTCTATATCATGGCAGATAAACAGAAGAATGGC
ATCAAAGTCAATTTTAAGATCCGGCATAATATCGAGGACGGATCAGTACAGCTTGCCG
ACCATTACCAACAGAATACACCGATAGGAGACGGTCCCGTCCTCTTGCCAGATAATCA

CTATTTGAGCACACAATCCGCTCTTTCTAAAGATCCGAATGAAAAGCGCGATCATATG
GTCTTGAAAGAGTTCGTCACTGCCGCCGGCATAACCGGCGGATCCGCCGGAAGCGGA
**gctactaacttcagcctgctgaagcaggctggagacgtggaggagaaccctggacct**ggcaccggttccaacatggcgattatcaag
gaattcatgaggttcaaagttcacatggagggatctgtaaacggccacgagttcgaaatcgagggagagggggagggtcgaccatatgaa
ggaacacagaccgctaaattgaaggttactaaaggtggaccacttcctttcgcatgggacattcttagcccccagtttatgtatggttccaagg
cgtatgtcaaacatcccgcagacattcctgattaccttaaactgagttttccagaggggttcaaatgggagcgcgtcatgaactttgaagatgg
gggcgttgttacagtgactcaggactcttctttgcaagatggagagtttatatataaggtgaaacttcgcggaaccaacttcccaagcgatggc
ccagtcatgcagaagaaaactatgggttgggaagccagtagcgagcggatgtacccagaggacggcgcacttaagggtgagattaagca
gcggctgaaactcaaagacggaggccattatgacgcggaagttaaaacgacgtacaaagctaaaaagcctgtccaattgccaggcgcgta
caatgtcaatattaaattggacattaccagtcacaacgaagattacacaatagtggagcaatacgaacgcgcagagggacgccattcaaca
gggtaa

**Hisx6-*Hy*MraY**

His tag is underlined. MraY gene is shown in pink.

CATCACCATCACCATCACATGATATACCATCTTGCCATACTCCTGAGGGAGCACTTCTT
CGCCTTCAACGTGCTCAAGTACATAACCTTCCGTTCTTTCACGGCAATACTCCTCGCC
TTCTTTATAACTCTGATACTTTCCCCCACCTTCATGAAGAAGTTTGCGAAGATACAGAG
ACTCTTCGGAGGGTACGTCAGAGAGTACACACCCGAACACCACGAGAGCAAGAAGT
ACACACCAACCATGGGCGGGGTTGTGATAGTAACGGTCATACTGATAACCTCAGTCCT
TCTCATGCGCCTTGATATCAGGTACACGTGGGTTCTCGTCTTTTCAACGCTGTCCTTTG
CCCTCATAGGGTTCGTGGACGACTGGATAAAACTCAAGAATAAAAAGGGTCTCTCAA
TAAAGGCGAAGCTCGCCTTCCAGATGTCCTTCGCTTTAGCCGTATCCCTGCTCATCTTT
TACTGGGTTGGTCTGGAGACAAAGCTATACTTTCCCTTCTTCAAGGAGCTCACCGTAG
ATCTGGGCTGGTTATATATACCCTTCTCAATGTTCATCATAGTAGGGACCGCTAACGCG
GTGAACCTTACCGACGGTCTGGACGGACTCGCCATAGGTCCTTCAATGACCACAGCA
ACAGCCTTCGGCGTGATAGCCTACGTGGTGGGTCACTCAAAGATAGCCCAGTACTTG
GGAGTCCCGCACGTTCCCTACGCAGGTGAGATAACGGTGTTCTGCTTCGCGATAATAG
GTGCAGGGCTTGGCTTTTTGTGGTTCAACACTTACCCGGCTCAGGTGTTCATGGGAG
ACGTGGGAGCTCTGGGTCTCGGGGCTGCCCTCGCCACGGTTTCAATTATGACCAAGT
CAGAGTTCCTGCTCGCCGTTGCCGGAGGTGTGTTCGTATTTGAGACGGTAACCGTGAT
ACTTCAGATAATCTACTTCAGAGCCACAGGAGGGAAGAGGCTCTTCAGAAAGGCTCC
CTTCCACCACCACCTTGAGGAGAAGGGGCTGGACGAACCCAAGATAGTGGTGAGGA
TGTGGATAGTTTCAGCCCTGCTCGCCATAGTGTCAGTGGCGATGCTGAAGCTCAGGTA
A

#### Supplementary Figures & Table

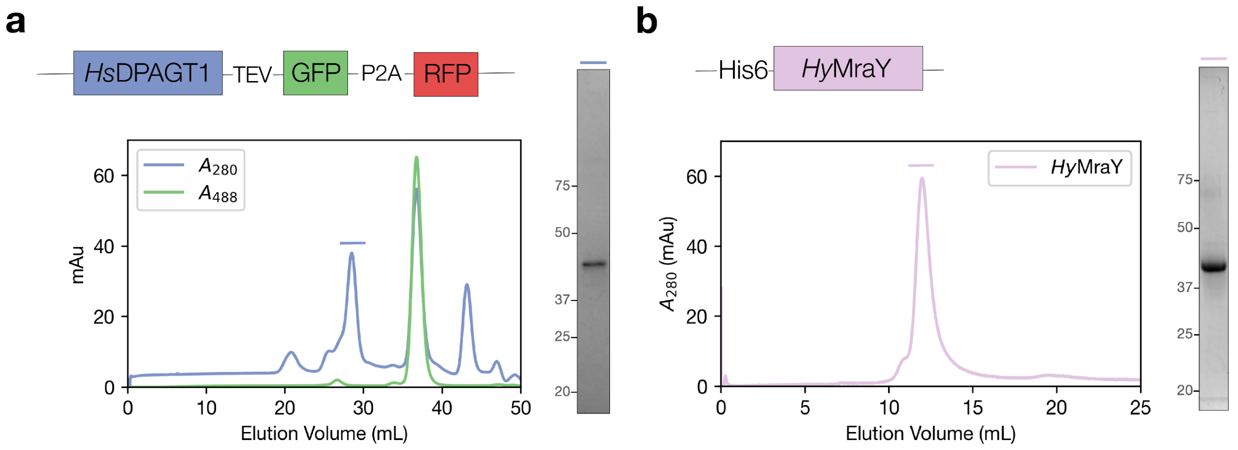


**Figure S1. Purification of *Hs*DPAGT1 and *Hy*MraY.** (a,b) Construct designs, representative size exclusion chromatography profiles, and SDS-PAGE gels of purified (a) DPAGT1 and (b) MraY. The indicated fractions from the chromatograms were used for grid preparation for cryo-EM. By comparing the 280 nm and 488 nm signals in (a), the indicated fraction is estimated to be 99 parts DPAGT1 to 1 part DPAGT1-GFP.

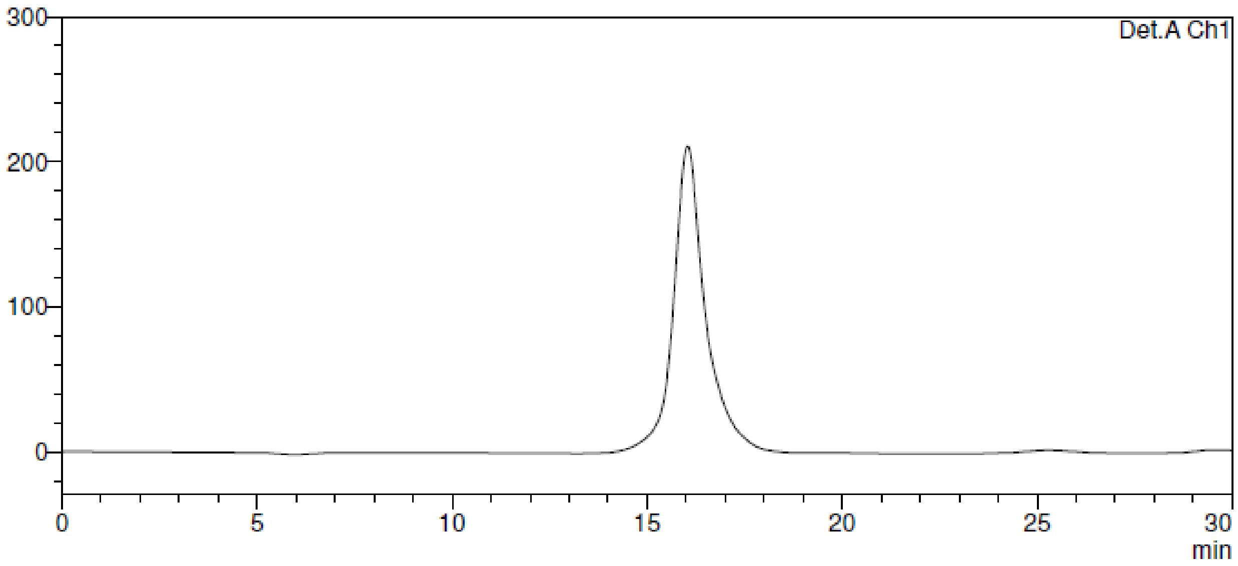


**Figure S2. HPLC analysis of APPB.** APPB (300 mg) was synthesized from uridine in 30% overall yield using a previously described synthetic protocol.[19,20] The purified compound was confirmed by HPLC analysis and was employed as the ligand material for cryo-EM experiments. HPLC conditions were: Phenomenex Kinetex 1.7 µ XB-C18 100 Å 150 x 2.10 mm column, 75:25 $CH_3OH$:0.05M $NH_4HCO_3$ aq, 1.0 mL/min flow rate, monitoring at 254 nm. The resulting sample had 98.8% purity.

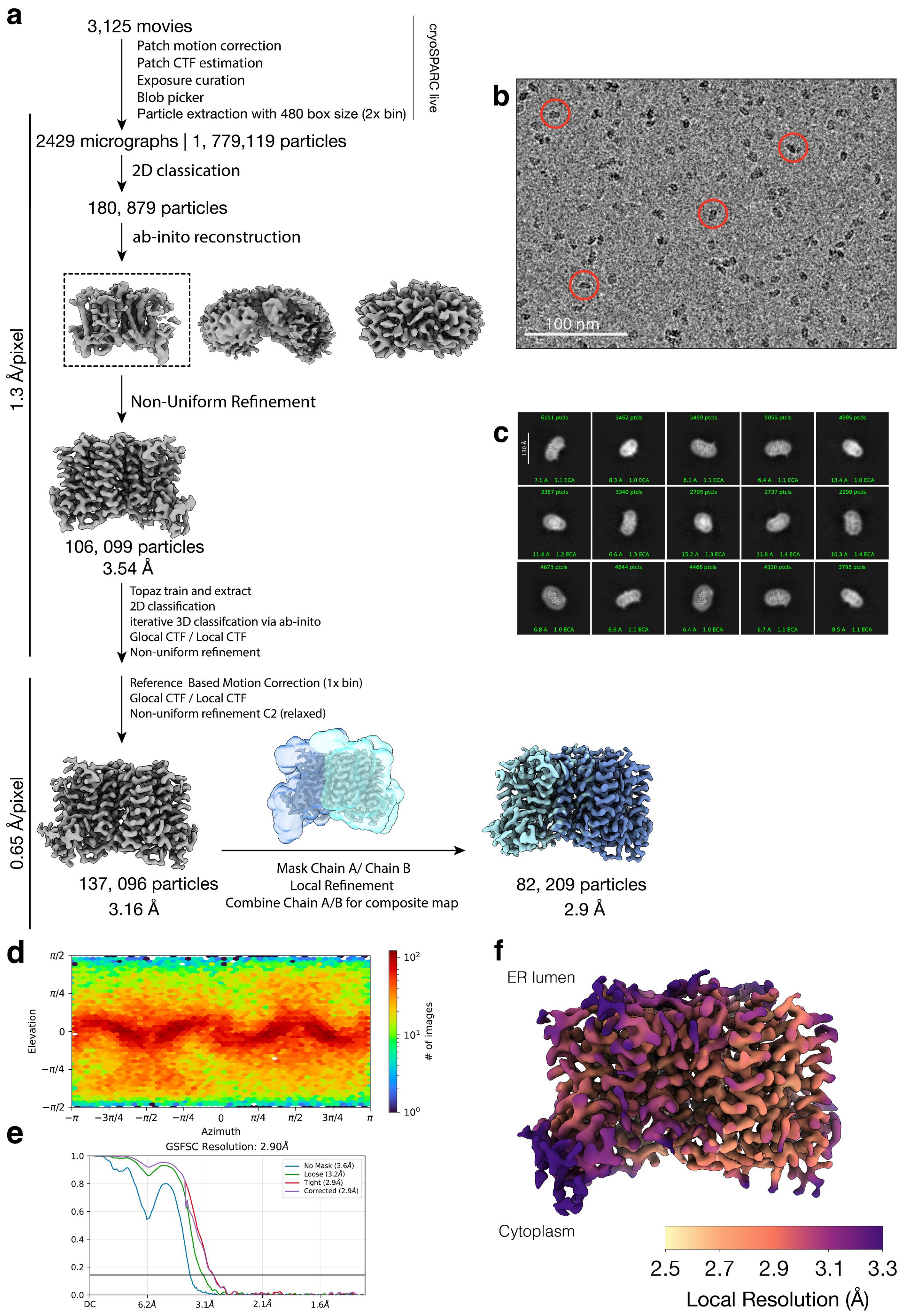

a
3,125 movies
Patch motion correction
Patch CTF estimation
Exposure curation
Blob picker
Particle extraction with 480 box size (2x bin)
cryoSPARC live
2429 micrographs | 1, 779,119 particles
2D classication
180, 879 particles
ab-inito reconstruction
Non-Uniform Refinement
106, 099 particles
3.54 Å
Topaz train and extract
2D classification
iterative 3D classifcation via ab-inito
Glocal CTF / Local CTF
Non-uniform refinement
Reference Based Motion Correction (1x bin)
Glocal CTF / Local CTF
Non-uniform refinement C2 (relaxed)
137, 096 particles
3.16 Å
Mask Chain A/ Chain B
Local Refinement
Combine Chain A/B for composite map
82, 209 particles
2.9 Å
1.3 Å/pixel
0.65 Å/pixel
b
100 nm
c
d
Elevation
Azimuth
### of images
e
GSFSC Resolution: 2.90Å
No Mask (3.6Å)
Loose (3.2Å)
Tight (2.9Å)
Corrected (2.9Å)
DC
6.2Å
3.1Å
2.1Å
1.6Å
f
ER lumen
Cytoplasm
2.5
2.7
2.9
3.1
3.3
Local Resolution (Å)

**Figure S3. Cryo-EM data processing and analysis for DPAGT1.** (a) Summary of cryo-EM data processing workflow for DPAGT1. (b,c) Representative micrograph (b) and 2D class averages (c). (d) Angular distribution of particles of the final 3D reconstruction. (e) Gold-standard Fourier shell correlation (GSFSC) curve. (f) Local resolution map of the transmembrane region in the cryo-EM composite map colored in inferno by resolution from 2.5 Å (yellow) to 3.3 Å (dark purple).

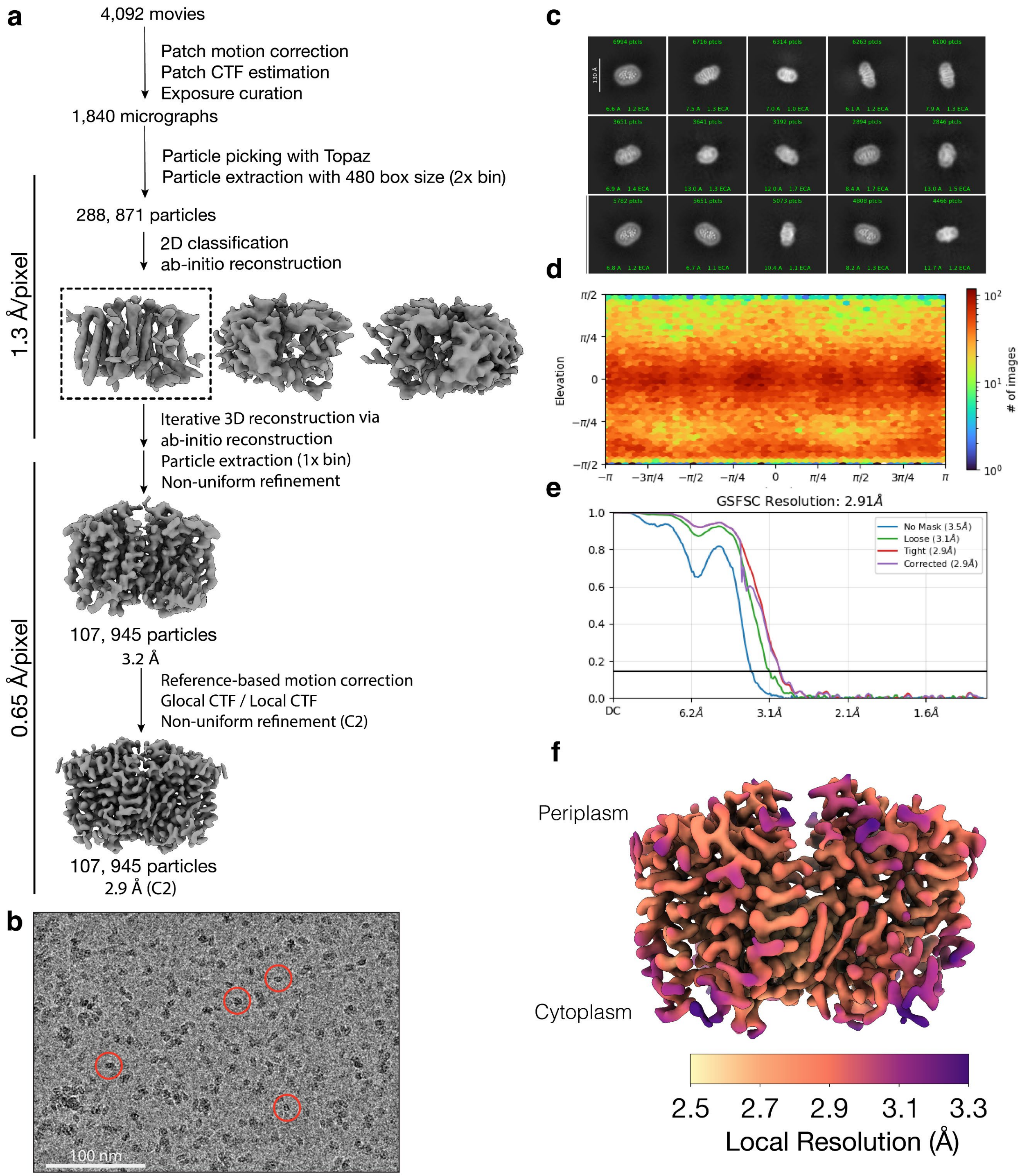


**Figure S4. Cryo-EM data processing and analysis for MraY.** (a) Summary of cryo-EM data processing workflow for MraY. (b,c) Representative micrograph (b) and 2D class averages (c). (d) Angular distribution of particles of the final 3D reconstruction. (e) Gold-standard Fourier shell correlation (GSFSC) curve. (f) Local resolution map of the transmembrane region in the final cryo-EM map colored in inferno by resolution from 2.5 Å (yellow) to 3.3 Å (dark purple).

**Table S1. Cryo-EM data collection, processing, refinement, and validation statistics.**

| | DPAGT1–APPB | MraY–APPB |
|---|---|---|
| PDB/EMDB Entry | 9ZNN/EMD-74451 | 9ZNO/EMD-74452 |
| **Data collection and processing statistics** | | |
| Microscope/Camera | FEI Titan Krios/Gatan K3 | FEI Titan Krios/Gatan K3 |
| Magnification | 130,000 | 130,000 |
| Voltage (kV) | 300 | 300 |
| Electron exposure ($e^{-}$/Å$^2$) | 70 | 70 |
| Defocus range (µm) | -1.0 to -3.0 | -1.0 to -3.0 |
| Raw pixel size (Å)<br>(Super-resolution mode) | 0.325 | 0.325 |
| Movies | 3,125 | 4,092 |
| Initial particle images (no.) | 1,799,119 | 288,871 |
| Final particle images (no.) | 82,209 | 107,945 |
| Symmetry imposed | C1 | C2 |
| Map resolution (Å)<br>(FSC threshold) | 2.9<br>(0.143) | 2.9<br>(0.143) |
| Map sharpening *B* factor (Å$^2$) | -79.7 | -109.8 |
| **Refinement and validation statistics** | | |
| Initial model used | *Hs*DPAGT1 AlphaFold2 model | *Hy*MraY AlphaFold2 model |
| Number of:<br>Non-hydrogen atoms<br>Protein residues<br>Ligands<br>Solvent molecules | <br>6,462<br>771<br>APB: 2; DPG: 2<br>80 | <br>5,436<br>662<br>APB: 2<br>0 |
| Min, max, mean *B* factors (Å$^2$):<br>Protein<br>Ligands<br>Solvent | <br>13.76, 170.99, 70.18<br>48.49, 129.04, 80.03<br>27.63, 109.35, 63.15 | <br>21.26, 147.43, 69.96<br>52.87, 103.57, 73.24 |
| R.m.s. deviations:<br>Bond lengths (Å)<br>Bond angles (°) | <br>0.004<br>0.655 | <br>0.003<br>0.589 |
| Ramachandran (%):<br>Favored<br>Allowed<br>Outliers | <br>98.69<br>1.31<br>0 | <br>99.08<br>0.92<br>0 |
| Poor rotamers (%) | 0 | 0 |
| Clashscore | 5.15 | 8.90 |
| MolProbity score | 1.27 | 1.48 |

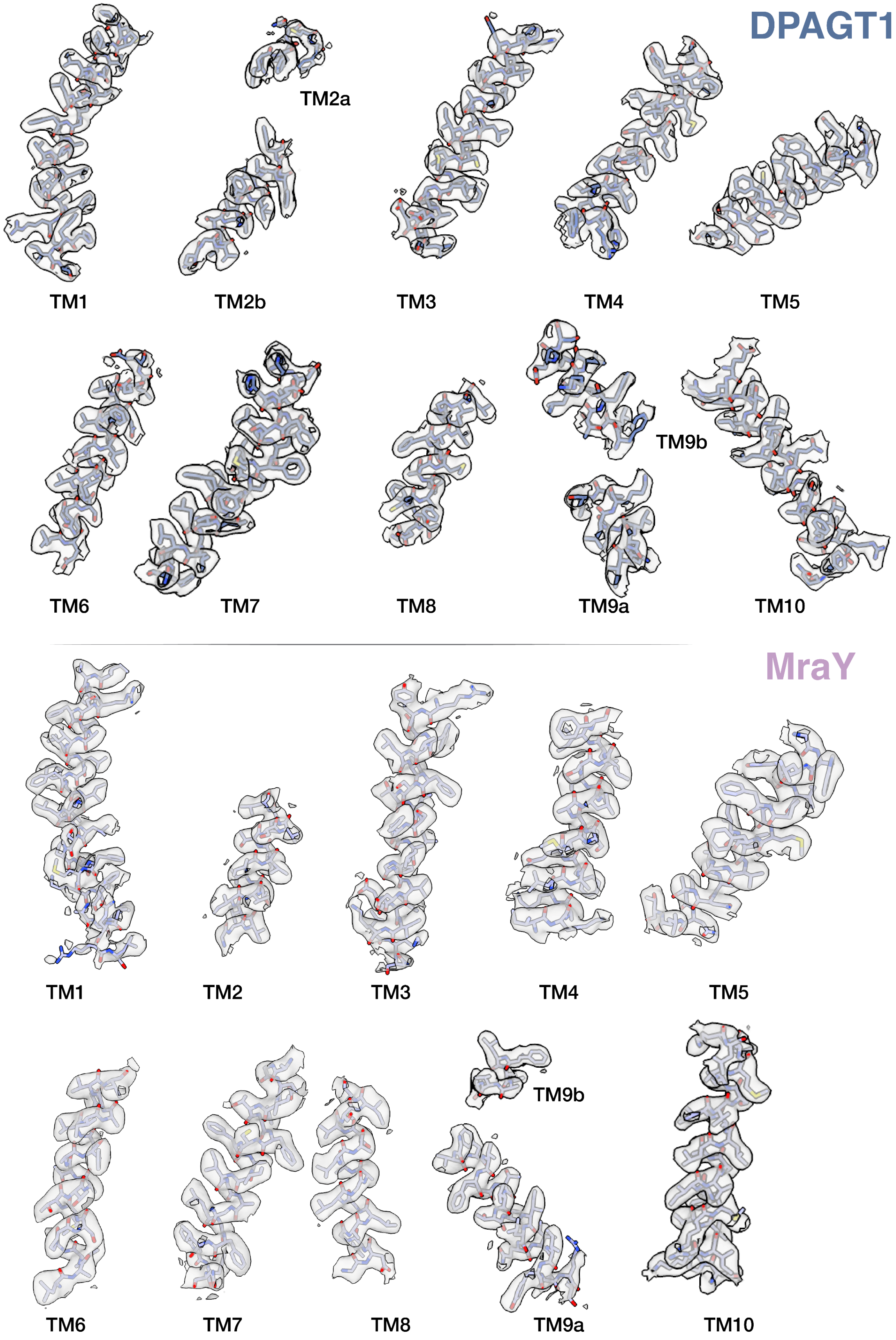
DPAGT1
TM1
TM2a
TM2b
TM3
TM4
TM5
TM6
TM7
TM8
TM9b
TM9a
TM10
MraY
TM1
TM2
TM3
TM4
TM5
TM6
TM7
TM8
TM9b
TM9a
TM10

**Figure S5. Model fits to cryo-EM density for transmembrane helices 1-10 in DPAGT1 and MraY.** All models are shown in sticks.

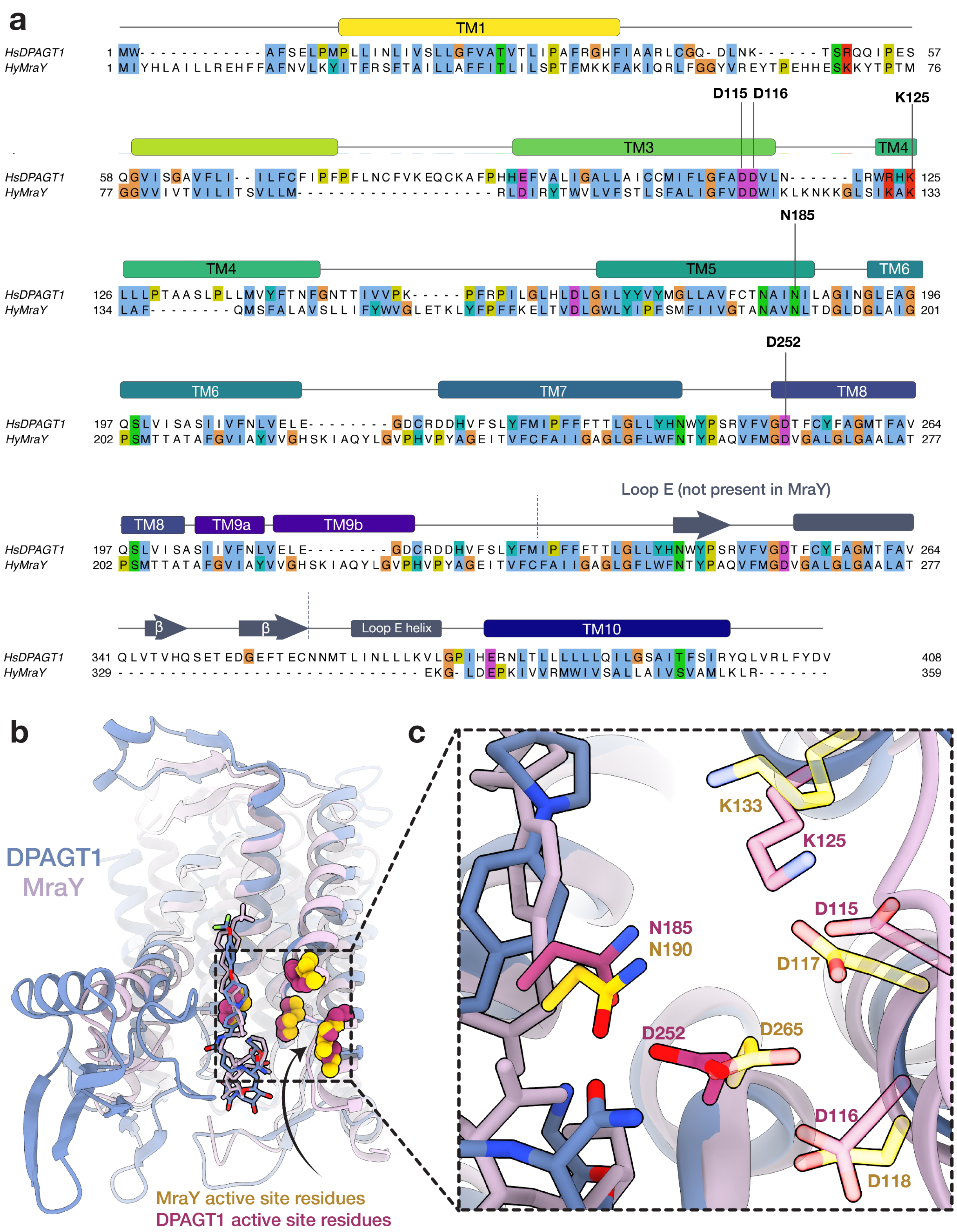


**Figure S6. DPAGT1 and MraY sequence and structural alignments.** (a) Structure-based sequence alignment using PROSMAL3D[21] with Clustal X[22] coloring for residues. Conserved active site residues are highlighted. Dashed lines demarcate Loop E insertion in DPAGT1 not present in MraY.[23] (b) Structural alignment of DPAGT1 (blue) and MraY (pink) with APPB *cis* conformer highlights the similar tertiary structures of the proteins. Dashed box indicates the active site, where conserved residues are shown as spheres in MraY (yellow) and DPAGT1 (purple). Mainchains are shown in ribbon form and APPB is shown in sticks. (c) Close up of boxed active site in (b). Sidechains of conserved residues are shown as sticks. Sidechains which are in contact with APPB are shown nontransparent, while residues which are not in contact are shown with 50% opacity. In $\textit{cis}_{\text{DPAGT1}}$, N185 and D252 contact APPB. In contrast, in $\textit{trans}_{\text{DPAGT1}}$, only D252 engages APPB (**Figure S11**). In both $\textit{cis}_{\text{MraY}}$ and $\textit{trans}_{\text{MraY}}$, N190 binds APPB (**Figure S11**). The residues which do not engage APPB – D115, D116, and K125 in DPAGT1 and D118, D119, K133, and D265 – may be targeted in future APPB iterations.

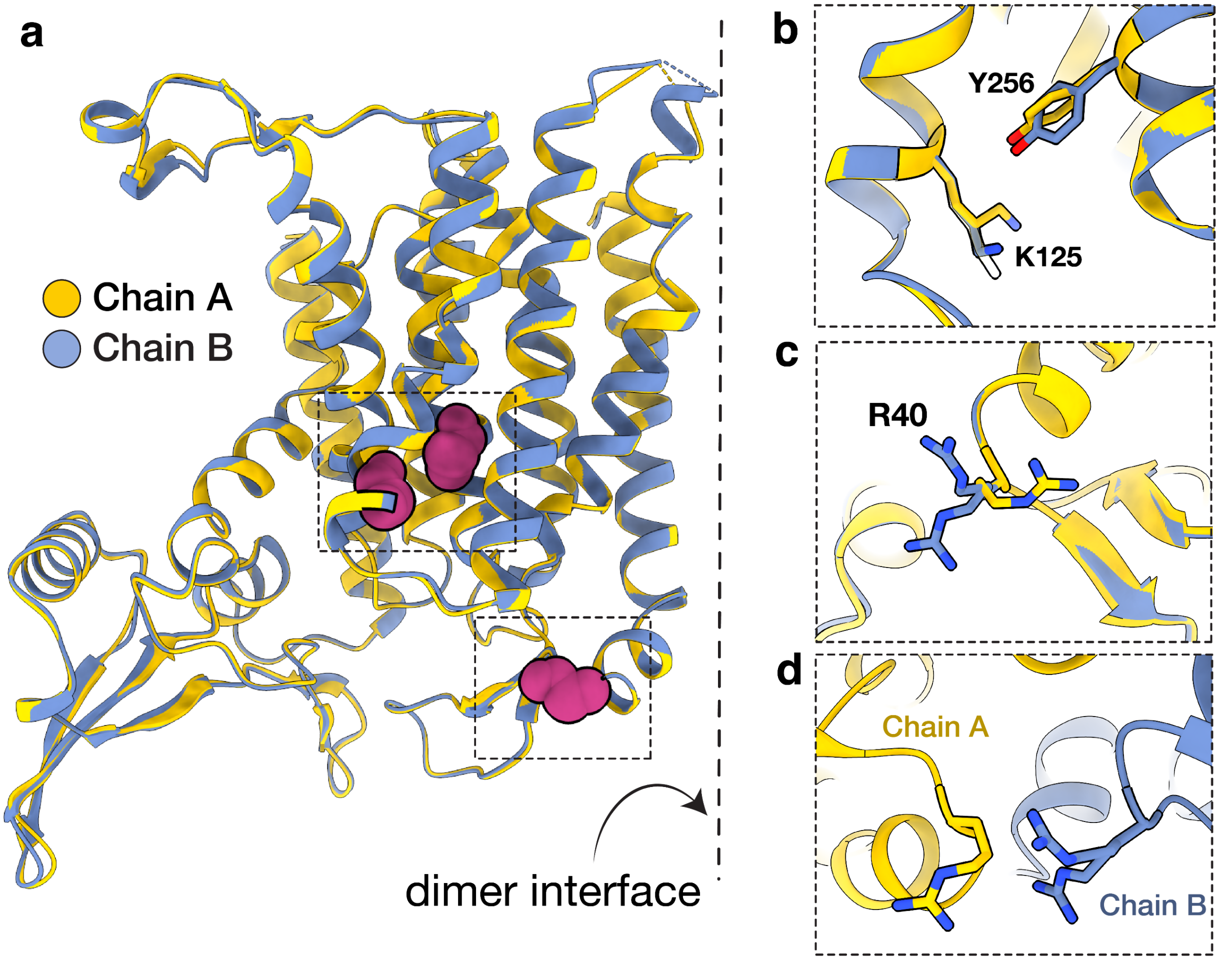


**Figure S7. DPAGT1 exhibits sidechain-level symmetry breaking.** (a) Structural alignment of DPAGT1 monomers shown as ribbons. The monomers possess nearly identical folds with an RMSD of 0.23 Å. However, at the residue level, several sidechains adopt different conformations between the monomers. Symmetry breaking residues are shown as spheres. Closeups of the residue sidechains are shown as sticks in (b) and (c). In (b), one K125 chain A conformer is silhouetted for clarity of the chain B residue. In (d), DPAGT1 dimer is shown to highlight symmetry breaking of R40 residues at the dimer interface.

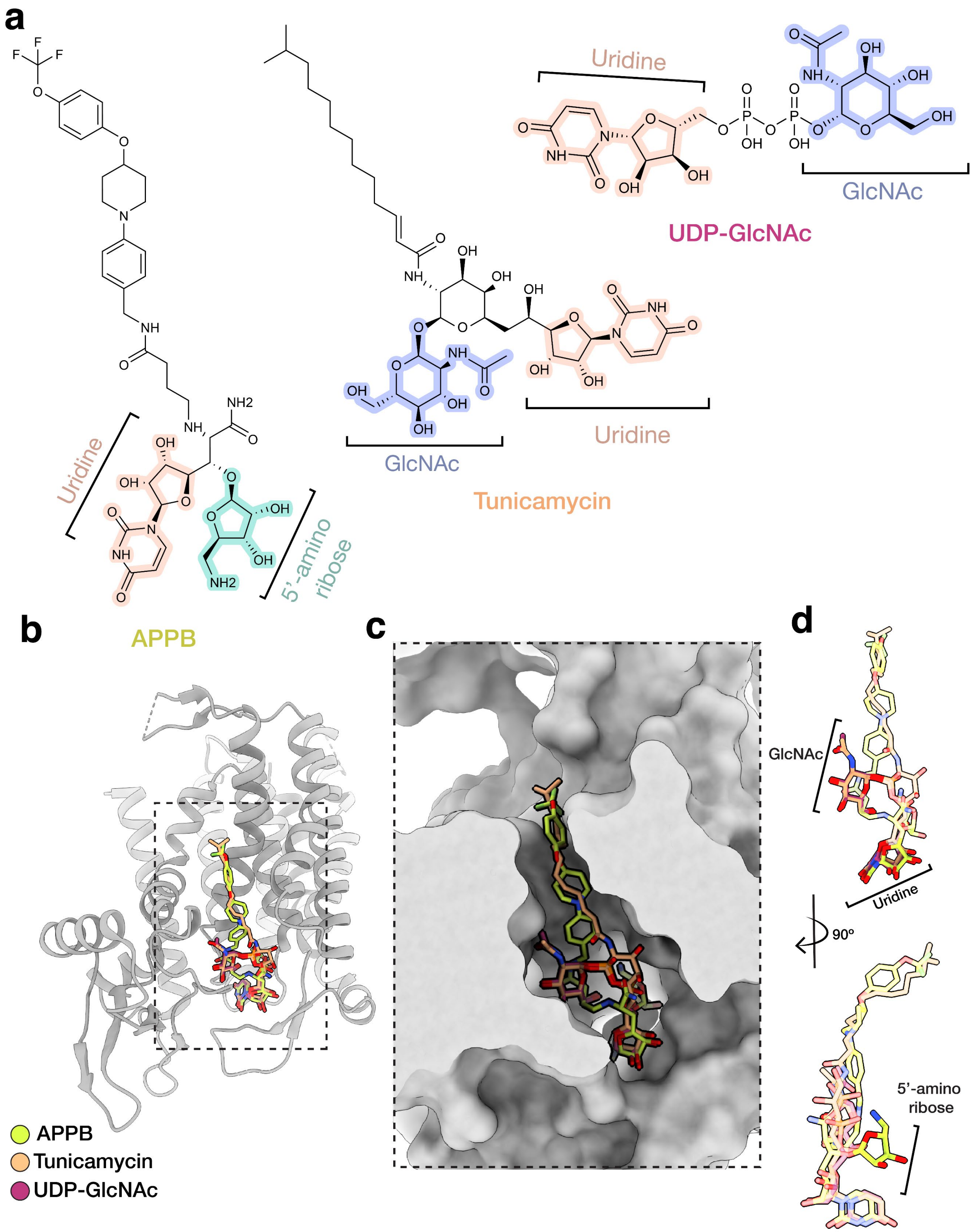


**Figure S8. DPAGT1 nucleoside ligands and ligand-bound structural comparisons.** (a) DPAGT1 inhibitors APPB and tunicamycin and substrate UDP-GlcNAc consist of uridine moieties (orange) and GlcNAc (blue) or 5'-aminoribose (turquoise) groups.[19,23,24] (b) Structural alignment of DPAGT1 with cis APPB (neon green), tunicamycin (orange), and UDP-GlcNAc (red) [PDBID in order: 9ZNN (this work), 5O5E[24], 6FWZ[24]]. Ligands are shown as sticks and protein is shown in ribbon form. (c) Molecular surface view of DPAGT1 binding groove with ligands. (d) Overlay of ligands. Uridine groups occupy highly similar positions. The APPB central amide carbonyl overlaps with the GlcNAc moieties in UDP-GlcNAc and tunicamycin.

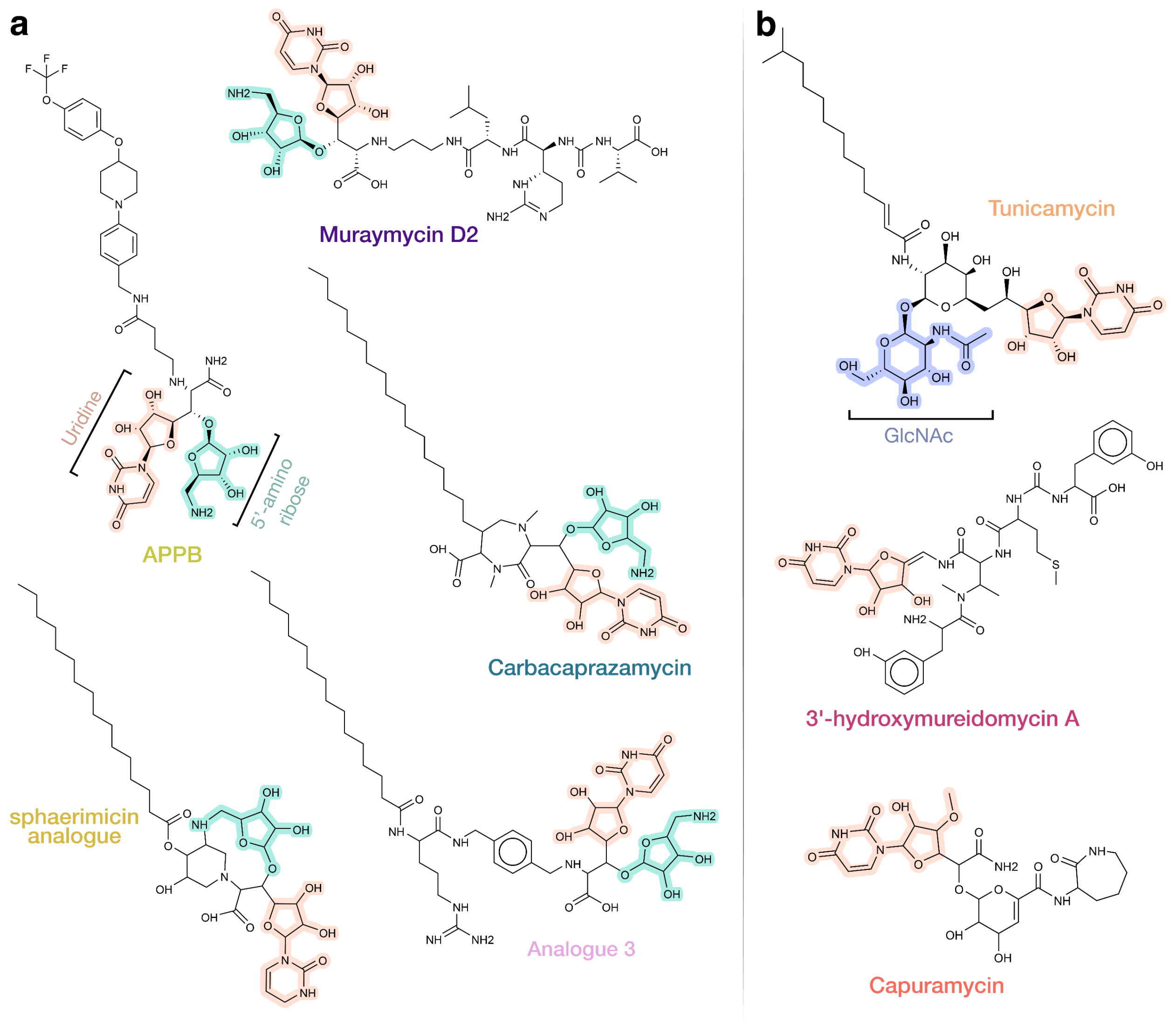


**Figure S9. Chemical structures of MraY nucleoside inhibitors.** All structurally characterized MraY inhibitors possess a uridine moiety (orange).[19,25–29] The inhibitors can be classified into those with (a) or without (b) 5’-aminoribose moieties (turquoise). Inhibitors lacking an aminoribose have only uridine or uridine and GlcNAc (blue) moieties.

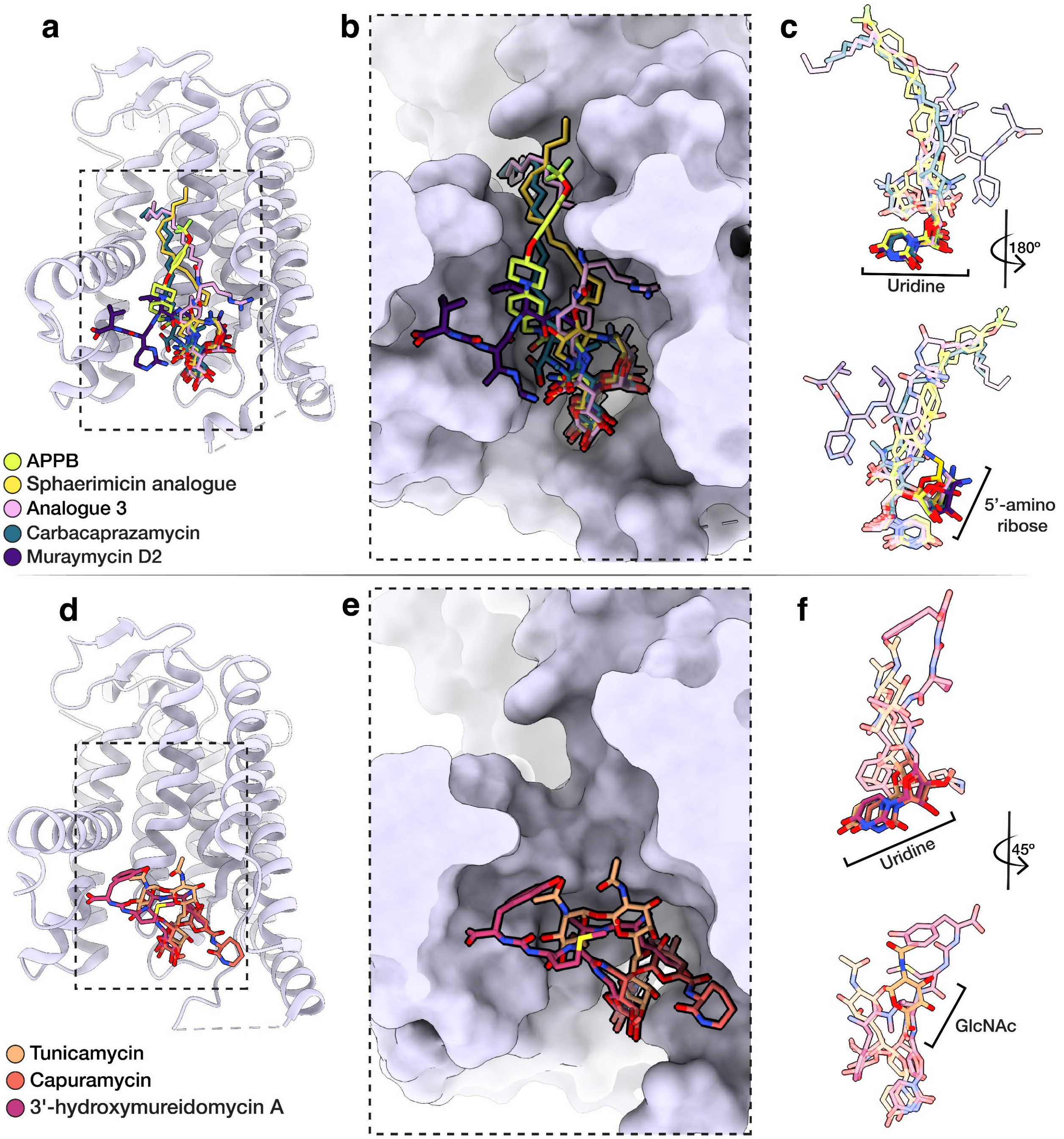


**Figure S10. Structural comparisons of MraY with inhibitors bound.** (a) Structural alignment of MraY and inhibitors with 5'-aminoribose moieties: APPB (neon green), sphaerimicin analogue (yellow), analogue 3 (pink), carbacaprazamycin (blue), muraymycin D2 (purple) [PDBID in order: 9ZNO (this work), 8CXR[27], 9B71[29], 6OYH[25], 5CKR[26]]. Inhibitors are shown as sticks and protein is shown in ribbon form. (b) Molecular surface view of MraY binding groove with inhibitors in (a) viewed from the cytoplasm. (c) Overlay of ligands in (a,b). The uridine and 5'-aminoribose moieties occupy the same position across the structures. (d) Structural alignment of MraY and inhibitors without 5'-aminoribose moieties: tunicamycin (light orange), capuramycin (orange), and 3’-hydroxymureidomycin (hot pink) [PDBID: 5JNQ[28], 6OZ6[25], 6OYZ[25]]. Note that the tunicamycin lipid tail is not shown because it was not modeled in ref. [28]. (e) Molecular surface view of MraY binding groove with inhibitors in (d) viewed from the cytoplasm. (f) Overlay of ligands in (d,e). The uridine moiety positions are highly analogous and occupy the same conserved uridine pocket in MraY.[30]

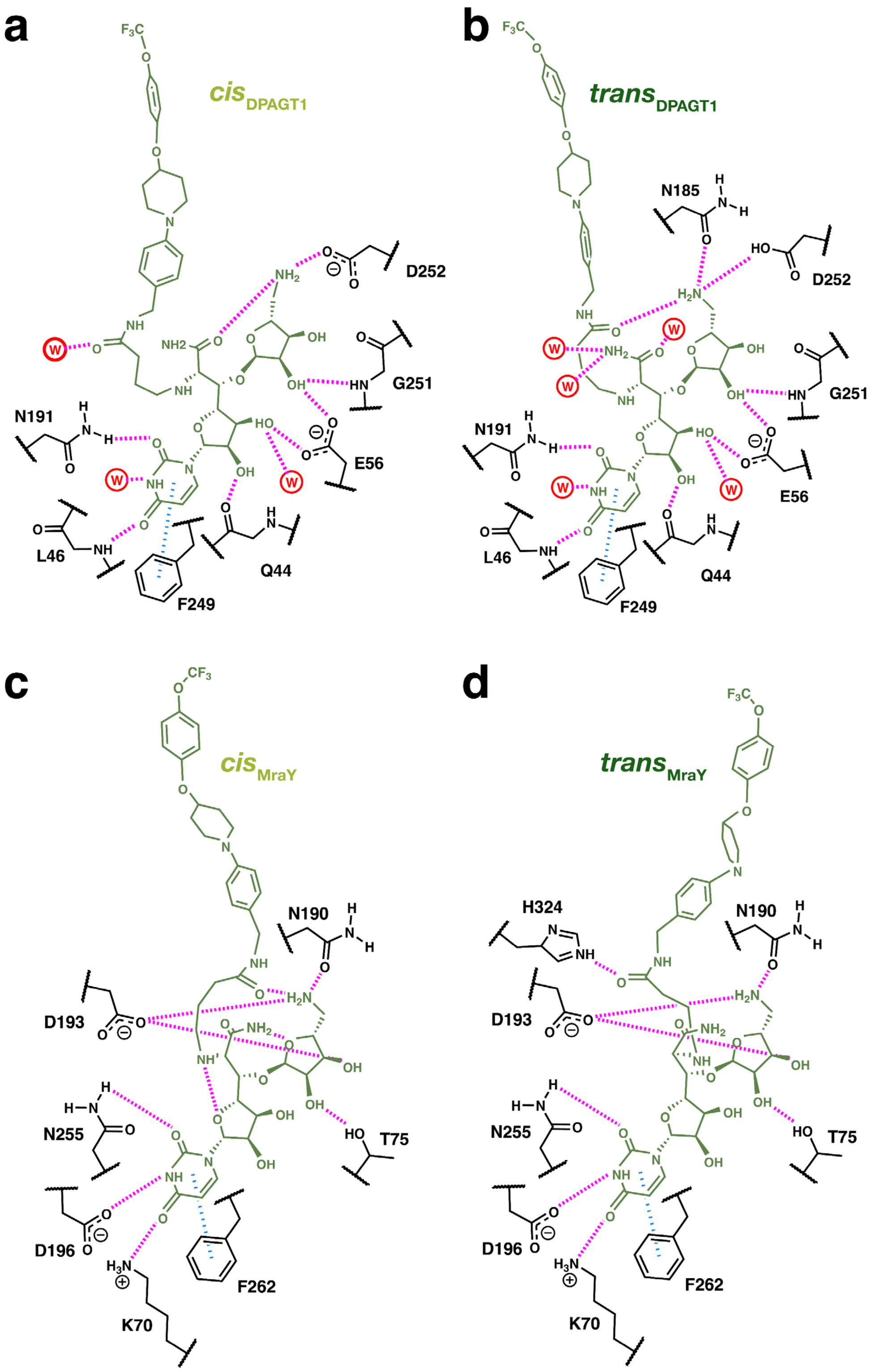


**Figure S11. Schematic representations of interactions made with APPB.** APPB contact maps for *cis*DPAGT1 (a), *trans*DPAGT1 (b), *cis*MraY (c), and *trans*MraY (d). APPB is shown in green, residues are shown in black, waters are shown in red, H-bonding interactions are shown in pink, and $\pi - \pi$ stacking interactions are shown in blue.

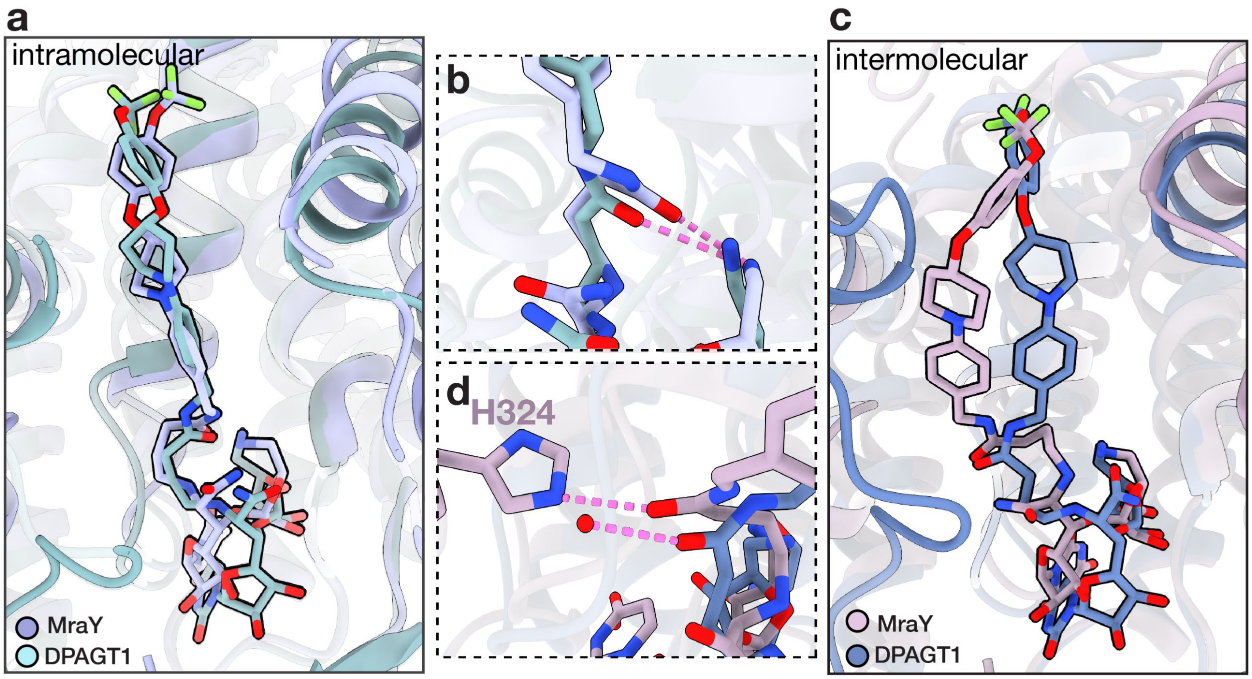


**Figure S12. Structural overlay of APPB conformers in DPAGT1 and MraY engaged in *intra*molecular or *inter*molecular H-bonds.** (a) The central amide carbonyl in *trans*$_{\mathrm{DPAGT1}}$ (blue) and *cis*$_{\mathrm{MraY}}$ (light purple) form intramolecular H-bonds, and APPB adopts highly similar geometries. APPB is shown as sticks and proteins are shown in ribbon form. (b) The carbonyls H-bond with the APPB aminoribose amines, as indicated by pink dashed lines. (c) The central amide carbonyl in *cis*$_{\mathrm{DPAGT1}}$ (blue) and *trans*$_{\mathrm{MraY}}$ (pink) form intermolecular H-bonds. The carbonyls adopt similar positions, but the TMPA lipid tails diverge in geometry. (d) The carbonyl H-bonds with a water molecule in DPAGT1 or H324 in MraY.

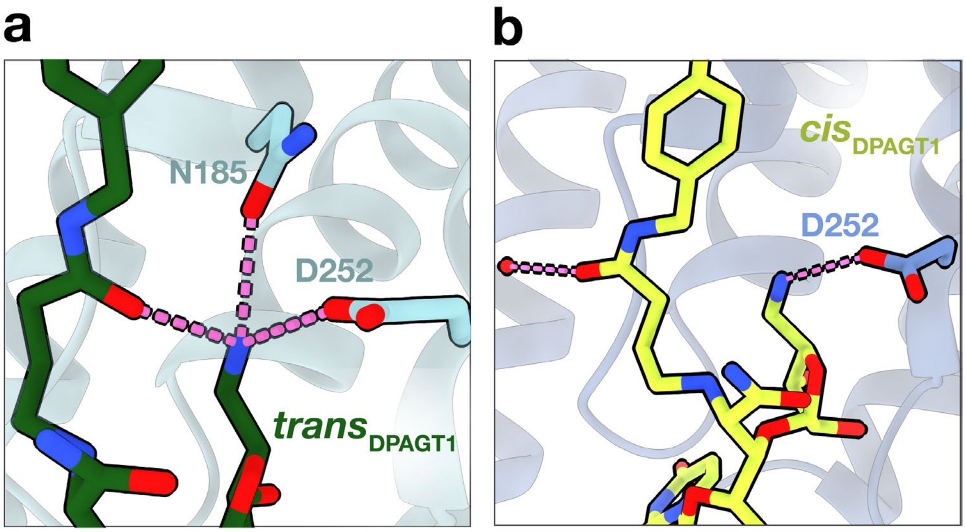


**Figure S13. H-bonds formed by APPB aminoribose amine.** In *trans*DPAGT1 (a), the amine forms three H-bonds with the central amide carbonyl and the N185 and D252 sidechains, thereby fulfilling its H-bonding potential. In contrast, in *cis*DPAGT1 (b), the amine H-bonds with the D252 sidechain.
